# Identification of stem cell marker-positive subpopulations in the vocal fold of the larynx through transcriptomic analyses

**DOI:** 10.1101/2025.03.25.645195

**Authors:** Keiichi Tamura, Hiroe Ohnishi, Koki Hasebe, Shintaro Fujimura, Tatsuya Katsuno, Zhaonan Zou, Shinya Oki, Yasuyuki Ohkawa, Koichi Omori

**Affiliations:** Department of Otolaryngology-Head and Neck Surgery, Graduate School of Medicine, Kyoto University, Kyoto, Japan; Center of Anatomical studies, Graduate School of Medicine, Kyoto University, Kyoto, Japan; Institute of Resource Development and Analysis, Kumamoto University, Kumamoto 860-0811, Japan; Division of Transcriptomics, Medical Institute of Bioregulation, Kyushu University, Fukuoka, Japan; Department of Head and Neck Oncology and Innovative Treatment, Graduate School of Medicine, Kyoto University, Kyoto, Japan

**Author notes:** Corresponding Author: Koichi Omori, Department of Otolaryngology-Head and Neck Surgery Graduate School of Medicine, Kyoto University, 54 Shogoin Kawahara-cho, Sakyo-ku, Kyoto City Kyoto 606-8507, Japan.

**Keywords:** laryngeal mucosa, scRNA-seq, spatial transcriptomics, laryngeal organoids, vocal fold stem cells, SOX9-positive basal cells, *Lgr5*-positive fibroblasts

## Abstract

Information on the maintenance of tissue homeostasis is important for the development of effective therapeutic methods, however, reports on the cellular composition and tissue stem cells of the larynx were scarce. Therefore, we analyzed mouse laryngeal mucosa using scRNA-seq, and spatial transcriptomics by photo-isolation chemistry, and performed the generation of laryngeal organoid used as an in vitro model from mouse laryngeal mucosa. As a result, we found a SOX9-positive basal cell subpopulation and a *Lgr5*-positive fibroblast subpopulation in the mouse vocal fold, and obtained three types of epithelial organoids from laryngeal epithelium. We also confirmed the differences in pseudostratified ciliated columnar epithelium composition between the supraglottis and subglottis of the mouse larynx. These findings provide novel insights and valuable tools for future research in laryngology and stem cell biology.

## Introduction

The larynx, located between the pharynx and trachea, serves essential functions such as breathing, airway protection, swallowing, and vocalization (1). Laryngeal dysfunction can lead to symptoms including chronic cough, aspiration, and voice disorders (1). Although regenerative therapies using biomaterials (2) and stem cells (3) have been explored for these conditions, their efficacy remains insufficient. Understanding the mechanisms that maintain tissue homeostasis is crucial for developing effective therapies; however, knowledge in this area is still limited for the larynx.

Various animal species have been used to evaluate vocal fold (VF) function (4, 5). While studies suggest that the human larynx responds to inhalation toxicity similarly to that of dogs and monkeys (6), mice are widely used in basic research due to their extensive genetic modification tools and molecular biology applications. The larynx consists of three anatomical regions: the supraglottis (SPG), glottis, and subglottis (SBG). In adult humans, the SPG comprises stratified squamous epithelium (SSE) and pseudostratified ciliated columnar epithelium (PCCE); the glottis consists of SSE, and the SBG is covered with PCCE (7). Similarly, in adult mice, the SPG and SBG contain SSE and PCCE, while the majority of the glottis consists of SSE (8). A notable anatomical difference is that only mice possess a laryngeal pouch, which may be necessary for ultrasonic vocalization (9). VFs have a layered structure comprising epithelium, lamina propria, and muscle. Human VF epithelium typically consists of 5–10 layers (10), whereas the mouse VF epithelium consists of three or four layers (8). Despite these structural differences, the SSE of the VF in both species expresses *Trp63* in basal cells, *Mki67* in cycling basal cells, and *Keratin13* (*Krt13*) in differentiated cells (8, 11). These similarities suggest that the mouse is a valuable model for investigating the mechanisms that regulate laryngeal mucosal homeostasis.

Recent advancements in scRNA-seq have facilitated research on tissue-specific cell populations across various organs. Studies on the larynx have also been conducted, but most focus on squamous cell carcinoma (12, 13). A single-cell atlas of the normal adult mouse larynx has been published; however, its primary objective was to examine microbial profile changes due to the rearing environment and their impact on immune-related cells rather than investigating laryngeal stem cells (14). In humans, the macula flava in the lamina propria has been identified as a potential VF stem cell niche (15). Additionally, ABCG2-positive side population cells, thought to include stem cells, have been detected throughout the VF (16). However, stem cells in the mouse laryngeal lamina propria remain poorly understood. Due to the narrow structure and sparse cell density of the mouse laryngeal lamina propria, gene expression analysis of specific regions using bulk RNA-seq is challenging. Recently, spatial transcriptomics method named photo-isolation chemistry (17) has been developed. This method enables gene expression analysis of only UV-irradiated small area, has been used for analysis in several research (18–20), and was also considered to be useful in the larynx. Furthermore, because the larynx is an organ with complex tissues and it is difficult to obtain sufficient samples for in vitro experiments, an appropriate in vitro model is required. Organoids of various tissues have been generated as organ models to investigate events occurring in vivo (21), but there have been no reports of larynx organoids Therefore, in this study, we aimed to elucidate the mechanisms that maintain laryngeal mucosal homeostasis, focusing on tissue stem cells. We analyzed laryngeal mucosa using scRNA-seq and spatial transcriptomics by photo-isolation chemistry and generated laryngeal organoids.

## Results

### Single-cell RNA-seq analysis of mouse laryngeal mucosa

The mouse larynx is divided into the SPG, glottis (including the VF), and SBG. The mucosal epithelium covering these regions consists of PCCE containing acetylated α-Tubulin (aTub)-positive ciliated epithelium and KRT13-positive SSE (Fig. 1a, Supplementary Fig. 1a-d). Tissue from the region marked by a red line in the figure was enzymatically dissociated into single cells for analysis (Fig. 1b). Cluster analysis was performed, and the Uniform Manifold Approximation and Projection (UMAP) visualization of the mouse laryngeal mucosa dataset is presented (Fig. 1c). Epithelial cell clusters were identified based on *Cdh1* and *Epcam* expression, while fibroblast clusters were distinguished by vimentin (*Vim*) and *Mmp2* expression (Fig. 1d).

**Fig. 1.**
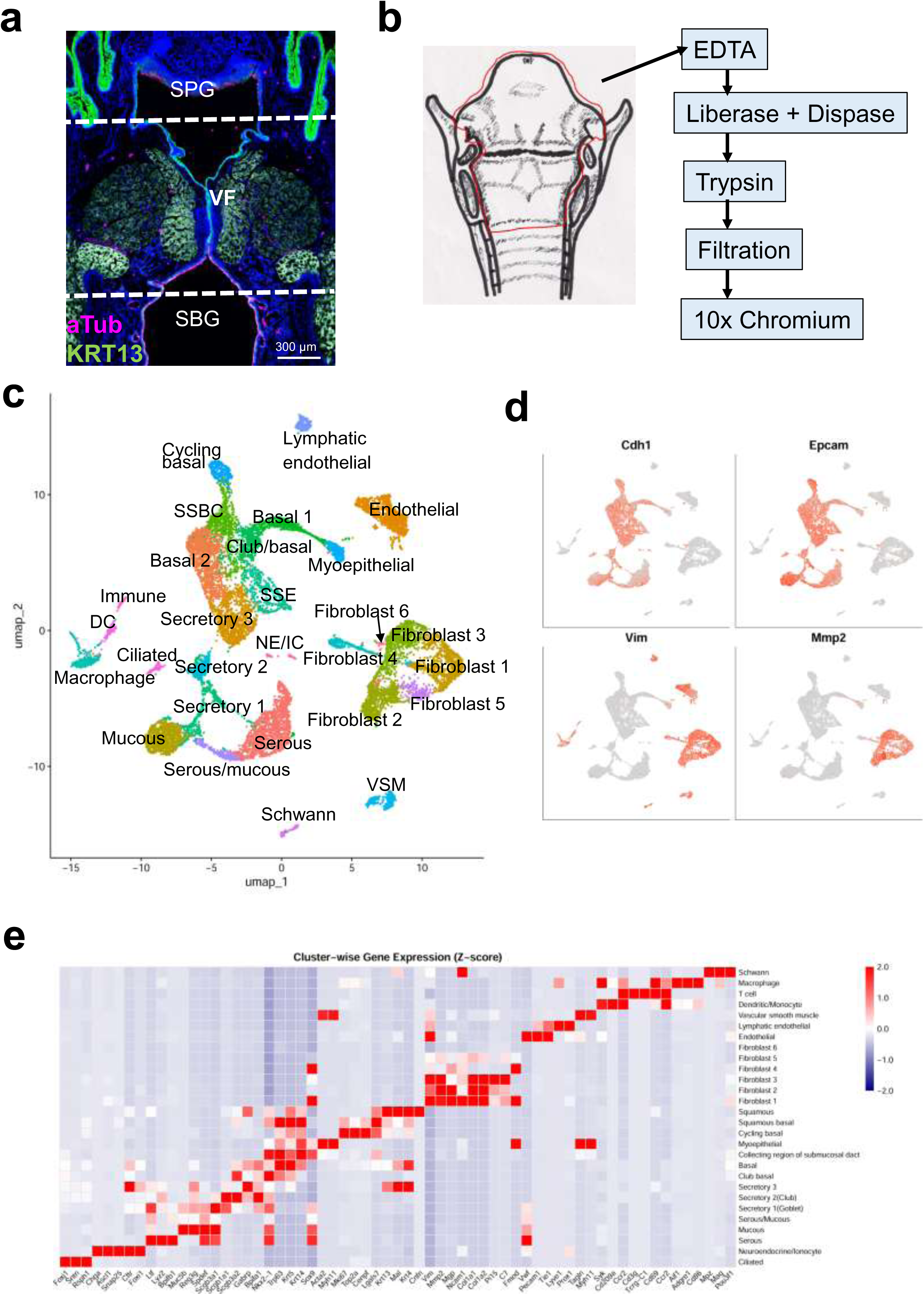
Single-cell RNA-seq analysis of the mouse laryngeal mucosa. **(a)** Representative immunofluorescence image of a coronal section of the mouse larynx, showing KRT13 (green), aTub (magenta), and DAPI (blue). Laryngeal regions are labeled with white letters and outlined with white dotted lines. Scale bar: 150 µm. (**b)** Schematic representation of the preparation method for a single-cell suspension of the mouse laryngeal mucosa. (**c)** UMAP visualization of the whole mouse laryngeal mucosa dataset. (**d)** Feature plot of *Cdh1*, *Epcam*, *Vim,* and *Mmp2*. (**e)** Heatmap of marker gene expression of each cell type. DAPI, 4′,6-diamidino-2-phenylindole; SPG, supraglottis; VF, vocal folds; SBG, subglottis; UMAP, Uniform Manifold Approximation and Projection; DC, dendritic cell; NE, neuroendocrine; SSE, stratified squamous epithelium; SSBC, stratified squamous basal cell; VSM, vascular smooth muscle; Cdh1, cadherin 1; Epcam, epithelial cell adhesion molecule; Vim, vimentin; Mmp2, matrix metalloproteinase-2.

Cell clusters were annotated based on differentially expressed genes (DEGs) and markers from previous studies. These included Basal cell 1 and 2 (*Krt5*, *Trp63*, *Krt14*), Cycling basal cell (*Mki67*, *Top2a*, *Cenpf*), Stratified squamous basal cell (SSBC) (*Trp63*, *Krt5*, *Krt14*, *Lgals7*), SSE (*Krt13*, *Mal*), Myoepithelial cell (*Acta2*, *Krt5*, *Myh11*), Club/basal cell *(Trp63*, *Nkx2-1*, *Scgb3a2*), Secretory cell 1, 2, and 3 (*Scgb3a2*, *Reg3g*, *Muc5b*, *Gabrp*, *Scgb1a1*), Serous cell (*Ltf*, *Lyz2*, *Nkx2-1*), Mucous cell (*Muc5b*, *Reg3g*, *Spdef*, *Nkx2-1*), Serous/mucous cell (*Ltf*, *Muc5b*, *Reg3g*, *Nkx2-1*), and Ciliated cell (*Foxj1*, *Sntn*) (Fig. 1c). A cluster expressing *Chga*, *Ascl1*, and *Snap25* was identified as the Neuroendocrine (NE) cell cluster. However, feature plots (Supplementary Fig. 2) of *Cftr* and *Foxi1* suggested that this cluster also includes Ionocyte (ICs) (Fig. 1c). Fibroblast subtypes (Fibroblast 1–6) were annotated based on *Vim* and *Mmp2* expression. Additional clusters included Endothelial cell (*Vwf*, *Pecam1*, *Tie1*), Lymphatic endothelial cell (*Pecam1*, *Prox1*, *Tie1*), Vascular smooth muscle cell (VSM) (*Acta2*, *Tagin*, *Myh11*), Dendritic cell/ monocyte (DC) (*Syk*, *Cd209a*), Immune cell (T cell; *Cd3g*, *Tcrg-C1*, *Cd69*), Macrophage (*Aif1*, *Adgre1*), and Schwann cells (*Mpz*, *Ncam1*) (Fig. 1c, Supplementary Fig. 2-1, 2, and 3). Expression levels of each marker gene are presented in the heatmap (Fig. 1e). These findings suggest that the cellular composition of the laryngeal PCCE closely resembles that of the tracheal PCCE and basal cells, secretory cells, SSE, and fibroblasts are divided into multiple clusters.

### Characterization of cell types in PCCE and submucosal gland epithelium clusters

The distribution of cells within each cluster of basal and secretory cells in the PCCE of the SPG and SBG was examined. The area marked by the white square in Figure 2a was defined as the SPG and SBG PCCE regions, and the images in Figure 2c were obtained from these regions. As shown in Figure 2b, the PCCE composes multiple cell types. The distribution of cells in each cluster was analyzed using the genes specifically expressed in each cluster, identified through feature plots as marker genes (Supplementary Fig. 2). Regarding secretory cell markers, *Muc5b* was expressed in Mucous, Secretory 1, Serous/mucous, and part of Secretory 2; *Reg3g* was expressed in Mucous, Secretory 1, Secretory 2, Serous/mucous, and part of Secretory 3; *Gabrp* was expressed in Secretory 3, Secretory 2, SSE, Basal 2, and part of Secretory 1; BPIFA1 was expressed in Basal 2, Secretory 1, Secretory 2, Secretory 3, and Club/basal cells. The distribution of *Muc5b*, *Reg3g*, and *Gabrp*-positive cells was examined by *in situ* hybridization (ISH), while BPIFA1-positive cells were analyzed using immunofluorescence (IF). *Muc5b*- and *Reg3g*-positive cells were observed in the luminal surface epithelia and submucosal glands (SMGs) of both the SPG and SBG, whereas *Gabrp*-positive cells were detected only in the luminal surface epithelia. Although BPIFA1-positive cells were present in the luminal surface epithelia and SMGs of both regions, their abundance was significantly lower in the SPG than in the SBG. Furthermore, most BPIFA1-positive cells in the SMG overlapped with the aTub-positive area, suggesting that BPIFA1-positive cells are enriched in the SMG duct’s collecting region. These findings suggest that Secretory 1 corresponds to secretory cells in the SMG, while Secretory 2 and Secretory 3 represent secretory cells in the luminal surface epithelia. A comparison of *Gabrp*- and BPIFA1-positive cells indicates that Secretory 3 comprises secretory cells in the SPG luminal surface epithelia, whereas Secretory 2 comprises secretory cells in the SBG luminal surface epithelia. Additionally, colocalization with the aTub signal suggests that Club/basal is the secretory cells in the SMG collecting region. The expression of KRT5 and KRT14 (known basal cell markers) and ACTA2 (a myoepithelial cell marker) was confirmed by IF (Fig. 2c). The abundance of KRT14-positive cells in the basal layers of the luminal surface epithelia in the SPG, as well as in the SMG of the SPG and SBG, suggests that Basal 1 represents SMG basal cells, while a portion of Basal 2 near the SSBC cluster corresponds to basal cells of the SPG luminal surface epithelia. These results indicate that distinct basal and secretory cell populations with different gene expressions are distributed across the luminal surface epithelia and SMG.

**Fig. 2.**
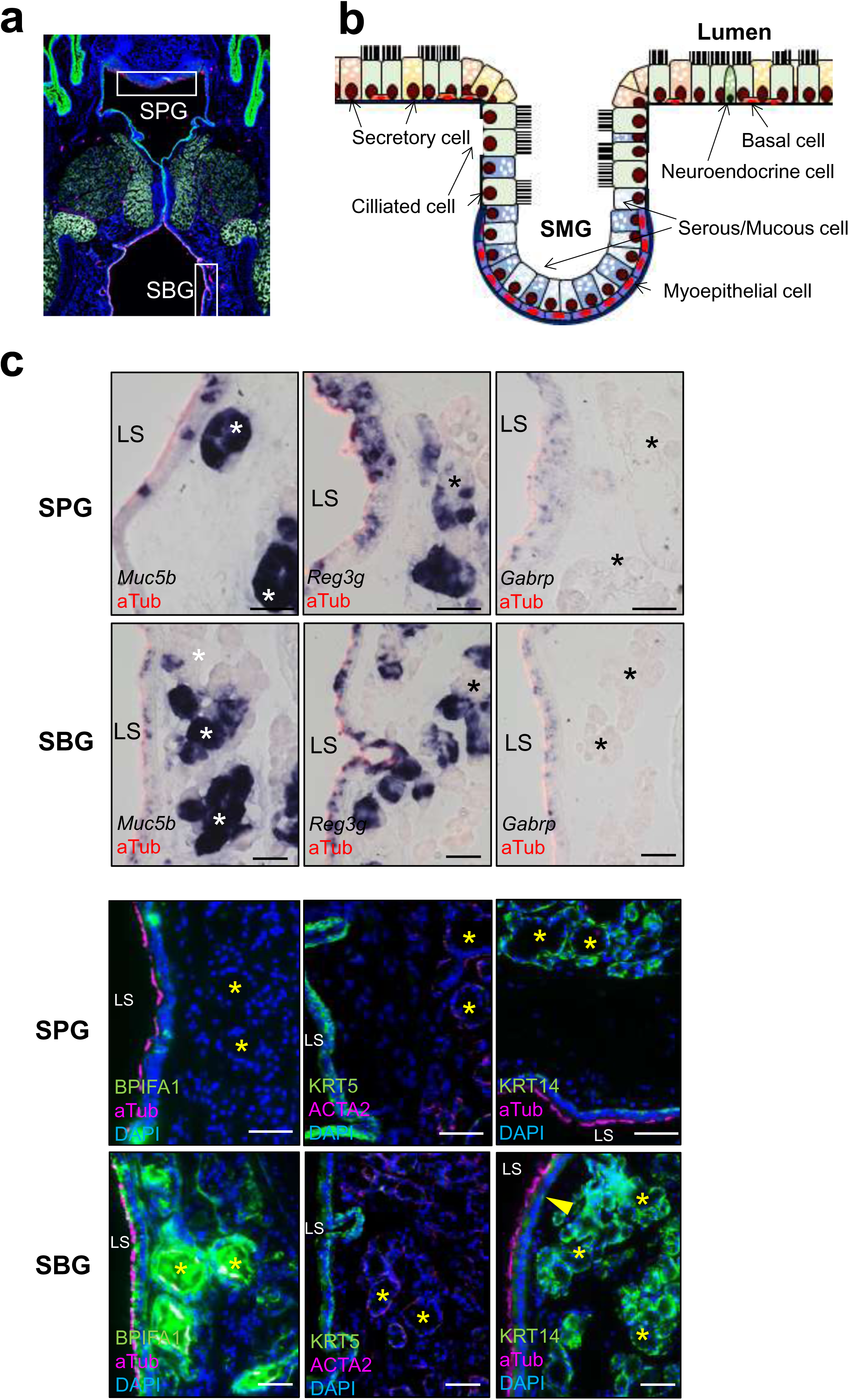
Characterization of gene expression in PCCEs of the laryngeal mucosa and SMG. **(a)** Coronal section of the mouse larynx showing the areas used as SPG and SBG in the following figures. **(b)** Schematic representation of the cell types constituting PCCEs and SMG epithelium. **(c)** Expression analysis of marker genes in SPG and SBG, including ISH images for *Muc5b*, *Reg3g,* and *Gabrp,* and IF images for aTub, BPIFA1, KRT5, ACTA2, and KRT14. Asterisks and arrowheads indicate SMG and KRT14 signal respectively. Scale bars: 50 µm. *n* = 3. SPG, supraglottis; VF,vocal folds; SBG, subglottis; LS, luminal surface; SMG, submucosal gland; PCCE, pseudostratified ciliated columnar epithelium; aTub, acetylated alpha-Tubulin; DAPI, 4′,6-diamidino-2-phenylindole.

### Characterization of Krt13-positive SSE clusters

Next, the SSE of the VFs was analyzed. Based on previous research (22), the VFs were divided into three regions: superior lateral, medial, and inferior lateral, with histological analyses performed on the medial region (Fig. 3a). The mouse VF epithelium is recognized as a thin SSE composed of several layers of basal and outer layer cells (Fig. 3b); however, its exact cellular composition has not been clarified. Therefore, *Krt13*-positive cells were extracted (Fig. 3c) and further subdivided into subclusters (Fig. 3d). In the mouse VF epithelium, basal layer cells express TRP63 (8), KRT13-strong positive cells localize near the surface, and LGALS7-, KRT14-, and MKI67-positive cells are predominantly found in the basal layer. Accordingly, each subcluster was annotated based on the feature plots for *Mki67*, *Trp63*, *Krt5*, *Krt14*, *Lgals7*, and *Krt13* in the UMAP of *Krt13*-positive cells (Fig. 3e, Supplementary Fig. 3). Since some Krt13-positive cells overlapped with secretory cell clusters, feature plots of *Reg3g* and *Bpifa1*, markers of secretory cell clusters, were also examined (Fig. 3e). These analyses suggested that subclusters 6, 3, 0, and 4 represent the PCCE lineage, while subclusters 6, 1, 2, and 5 correspond to the SSE lineage. The expression of SSE markers was further examined by IF. TRP63, KRT14, and LGALS7 were expressed in basal cells, whereas KRT13 was strongly expressed in the luminal surface layer (Fig. 3f–h). Array tomography analysis revealed that VF SSE consists of one to at least three cell layers along the dorsoventral axis (Supplementary Fig. 1e, movie). These findings suggest that the SSE in the medial VF composes 1–3 layers, which can be classified into four subclusters (6,1, and 2, 5), including *Mki67*-positive and *Mki67*-negative basal cell subclusters (6 and 1), based on gene expression.

**Fig. 3.**
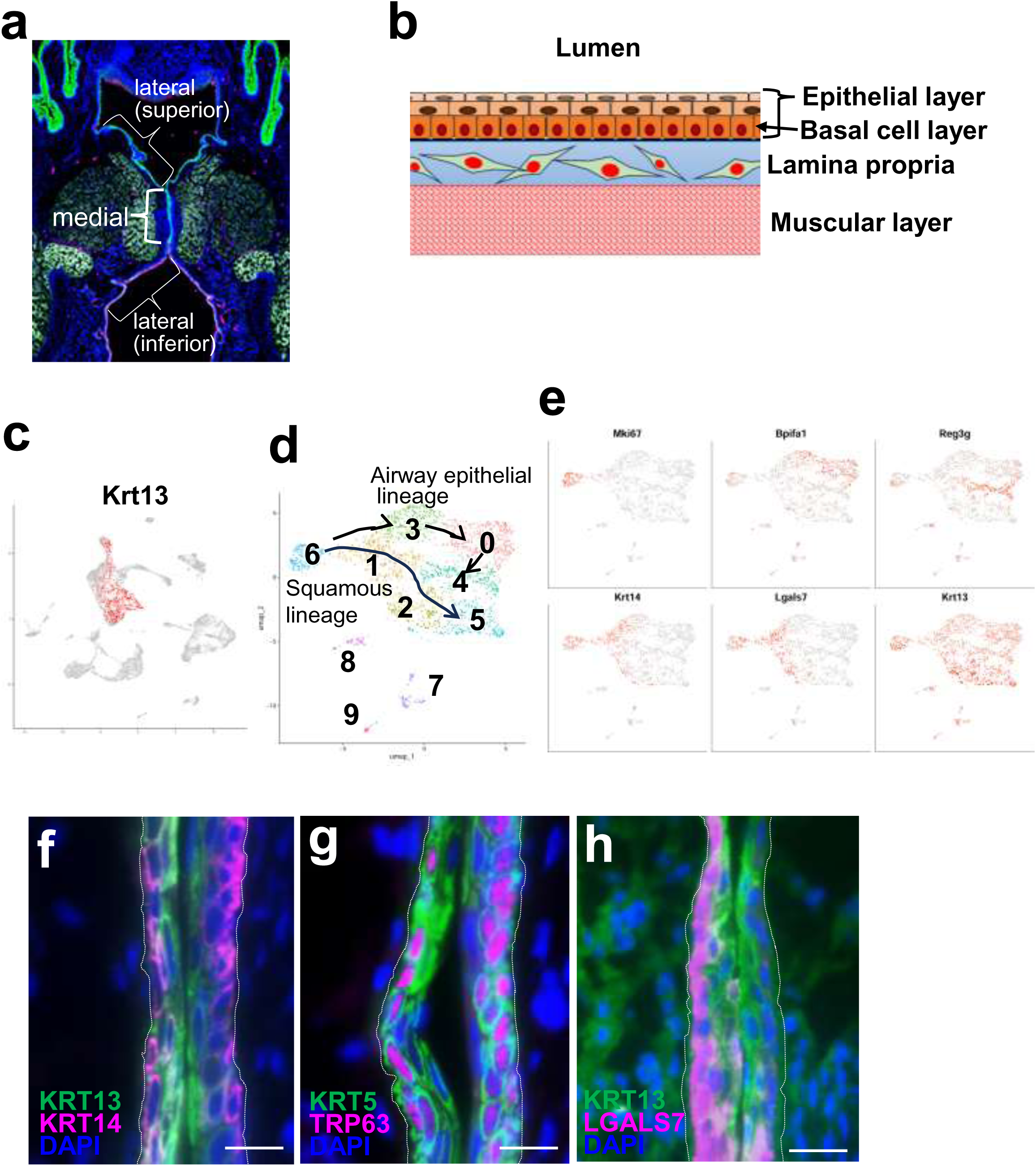
Characterization of Krt13-positive SSE clusters. **(a)** Coronal section of the mouse larynx showing each region of the VFs. (**b)** Schematic representation of SSE in the medial region of the VFs. **(c)** Feature plot of *Krt13*-positive cell data set used for subclustering in total UMAP. **(d)** UMAP visualization of the *Krt13*-positive cell dataset, confirming nine distinct clusters. **(e)** Feature plots of cycling cell marker, *Mki67*; basal cell markers *Krt14* and *Lgals7*; PCCE markers *Bpifa1* and *Reg3g*; and SSE marker *Krt13*. (**f-h)** IF images of the VF medial region in coronal sections for **(f)** KRT13 and KRT14, **(g)** KRT5 and TRP63, and **(h)** KRT13 and LGALS7. White dotted lines indicate the basement membrane location. Scale bars: 12.5 µm. *n* = 3. SSE, stratified squamous epithelium; UMAP, Uniform Manifold Approximation and Projection; DAPI, 4′,6-diamidino-2-phenylindole.

### Identification of Sox9-positive SSE stem cell subpopulation

Feature plots for *Sox9* showed *Sox9*-positive cells in both the Cycling basal cell and Basal cell clusters (Fig. 4a). The distribution of *Trp63* and *Lgals7* in the feature plot (Supplementary Fig. 2) suggested that these cells belonged to the basal cell region of the SSE. Similar to the feature plots for *Mki67*, *Sox9*, and *Trp63* (Supplementary Fig. 2 - 3), IF confirmed the presence of SOX9-positive cells among the TRP63-positive basal cells (Fig. 4b left), and some of them also overlapped with EdU-labeled cells (Fig. 4b right). Next, potential age-related differences in the distribution or number of SOX9-positive cells were investigated. Axial serial sections were prepared from the upper edge of the arytenoid cartilage to the closed portion of the VFs, and the number of SOX9-positive cells in the epithelium corresponding to the luminal surface of the laryngeal cavity was counted. Representative axial sections from young (left) and old (right) adult mice are shown in Fig. 4c. The median number of SOX9-positive cells in young and old adult mice was 7.5 (25–0) and 4 (16–0), respectively, while the number of EdU-labeled SOX9-positive cells in young and old adult mice was 3 (10–0) and 1 (9–0), respectively. These values demonstrated a significant decline with aging (Fig. 4d). These results indicate that SOX9-positive basal cells are present in the SSE from the superior lateral to the medial VF region, with some possessing proliferative potential.

**Fig. 4.**
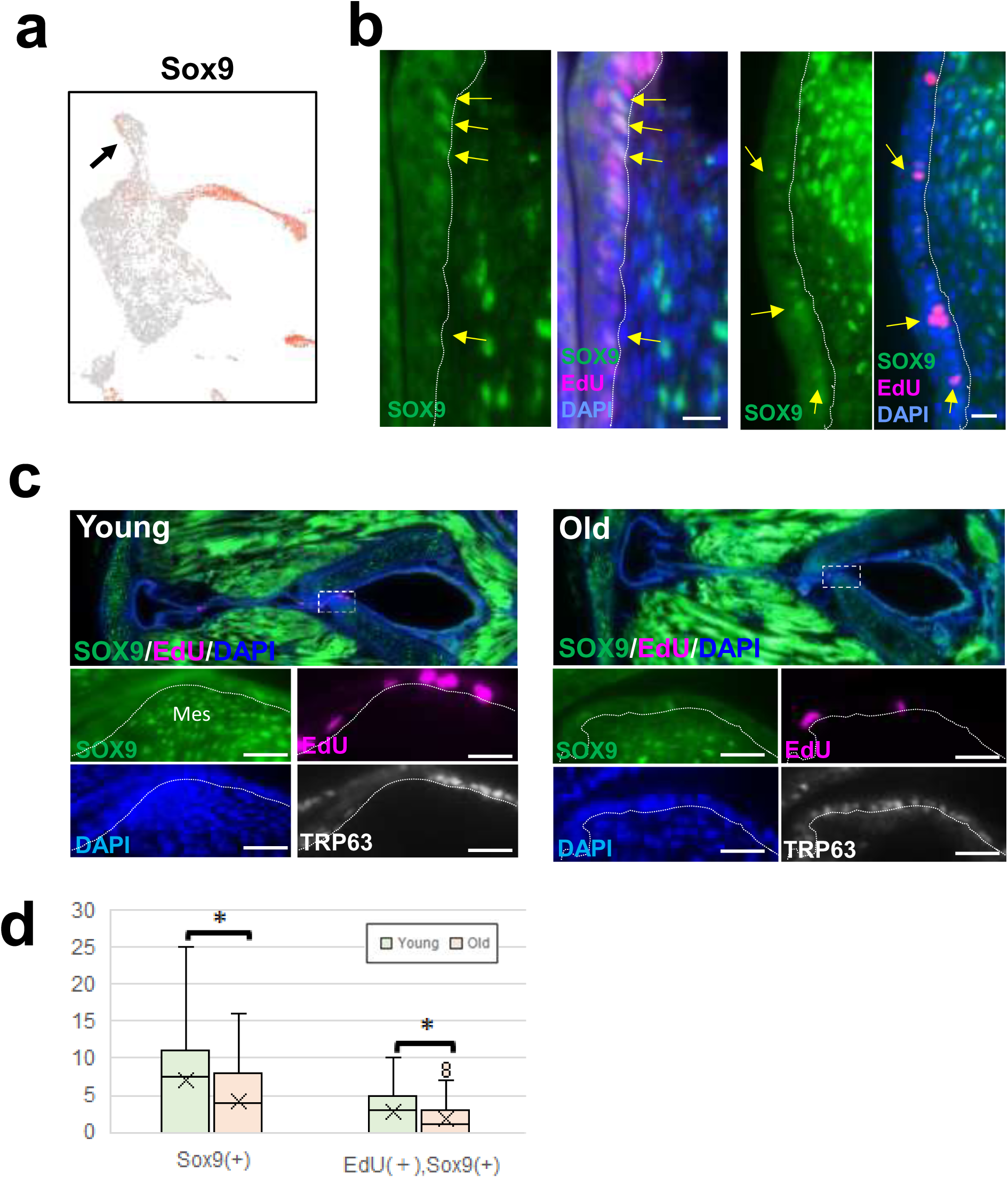
Identification of SOX9-positive SSE stem cell subpopulation. **(a)** *Sox9* expression in the cycling basal cluster in total UMAP (black arrow). (**b)** Representative images of SOX9-positive cells (green, indicated by yellow arrows) in the epithelia of VFs medial region. These cells represent a basal cell subpopulation, confirmed through co-staining with TRP63 (left, magenta) or EdU uptake (right, magenta). Scale bars: 12.5 µm. *n* = 3. (**c)** SOX9, P63, and EdU-positive cells in VFs of young and old adult mice. Representative IF images show SOX9 (green), TRP63 (white), and EdU (magenta)-positive cells in axial sections of VFs from young (left) and old (right) adult mice. Bottom images present high-magnification views of the white dotted squares in the top images. Scale bars: 25 µm. *n* = 3. (**d)** Quantification of SOX9-positive cells and SOX9-EdU double-positive cells in the VFs of young and old adult mice. Asterisks indicate significant differences (*p* < 0.05, Mann–Whitney U test). White dotted lines mark the basement membrane location. SSE, stratified squamous epithelium; EdU, 5-Ethynyl-2’-deoxyuridine; DAPI, 4′,6-diamidino-2-phenylindole.

### Examination of organoid formation ability of the cells derived from each region of mouse laryngeal mucosa

To validate the findings from the transcriptome analysis, organoids were generated as an in vitro model. Single cell suspensions were prepared from the laryngeal mucosa of the SPG, VF, and SBG, and organoids were generated by embedding culture in Matrigel (Fig. 5a). The organoids were morphologically classified into three types: Hollow, which were large and formed with a thin membrane; Solid, which were relatively small and formed with a thick membrane; and Tubular, which exhibited multiple branches (Fig. 5b). An analysis of each type organoid distribution across regions revealed the following median numbers: SPG Hollow 28 (48–14); SPG Solid 6 (9–1); SPG Tubular 16.5 (25–7); VF Hollow 23 (84–3); VF Solid 23.5 (43–13); VF Tubular 5 (10–0); SBG Hollow 51.5 (79–31); SBG Solid 10 (28–4); and SBG Tubular 12.5 (14–4). These data suggest that solid-type organoids were more prevalent in the VF, hollow-type organoids were more abundant in the SBG, and tubular-type organoids were more common in the SPG (Fig. 5c). Next, to determine whether laryngeal mucosal organoids contained PCCEs, SSEs, and SMG epithelia (SMGE), which constitute the larynx, we confirmed the expression of each epithelial marker by IF and ISH. These organoids exhibited a cystic structure with the apical surface facing inward (Fig. 5d). IF and ISH results identified three distinct organoid types derived from each mucosal region: BPIFA1, *Reg3g*, KRT14, and aTub-positive PCCE organoids; KRT13 and LGALS7-positive SSE organoids; and KRT5 and ACTA2-positive SMGE organoids (Figs. 5d and e). Similar to mouse VFs, SSE organoids also contained SOX9-positive cells in the basal layer (Fig. 5e). These findings indicate that organoids containing all three epithelial types were successfully generated from the laryngeal mucosa.

**Fig. 5.**
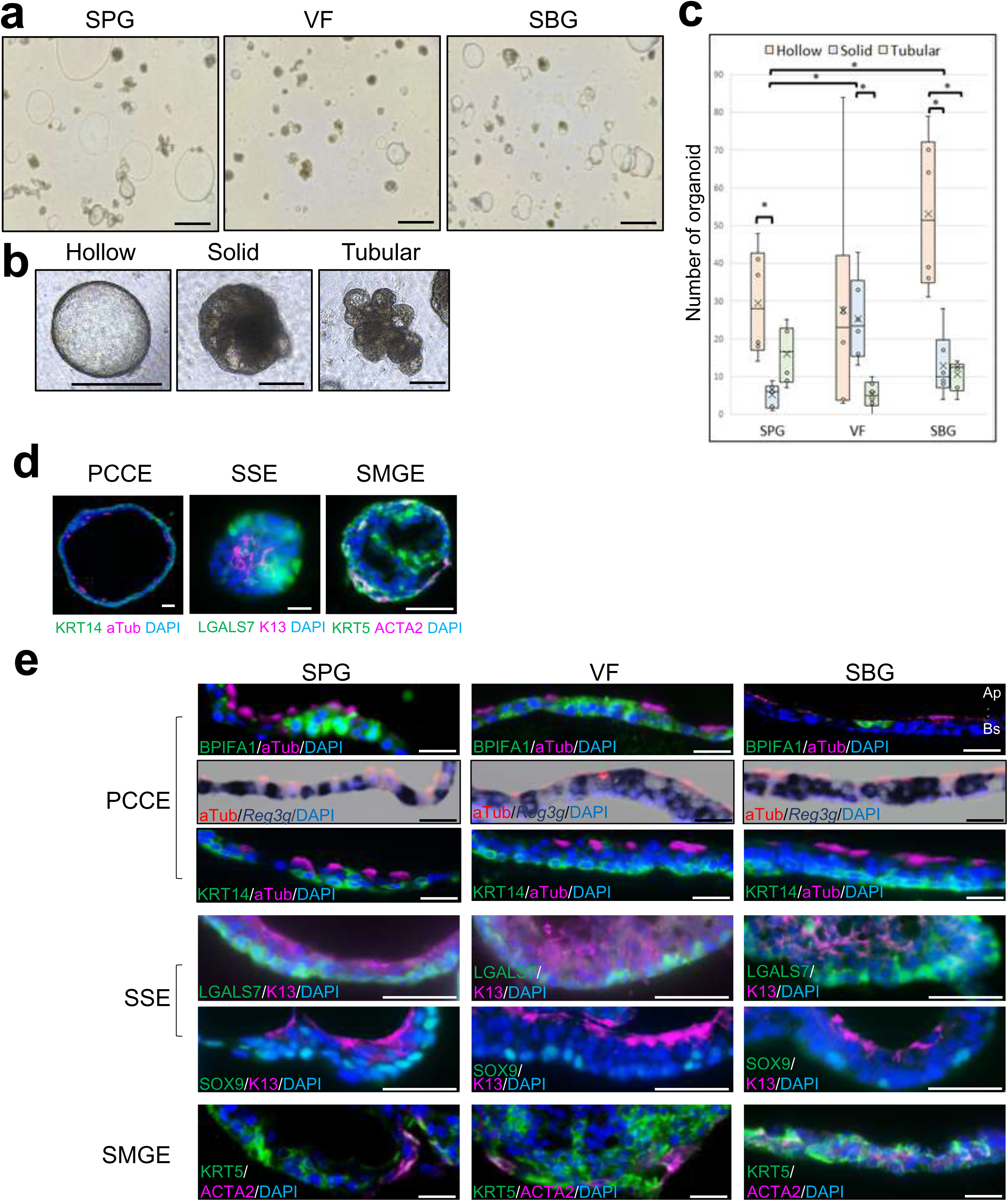
Examination of organoid formation ability in cells derived from different regions of the mouse laryngeal mucosa. **(a)** Representative images of organoids generated in Matrigel from each region of the laryngeal mucosa. Scale bars: 1 mm. **(b)** High-magnification images of three distinct types of organoids. Scale bars: 200 µm. **(c)** Quantification of organoids cultured from each laryngeal mucosal region. *n* = 6. Asterisks indicate significant differences (*p* < 0.05, Wilcoxon signed-rank sum test). **(d)** Expression of PCCE, SSE, and SMGE marker in organoids derived from different laryngeal mucosal regions. Low-magnification images of the three organoid types are shown: PCCE organoids contain KRT14 (green) and aTub(magenta) positive cells. SSE organoids contain LGALS7 (green) and KRT13 (magenta) positive cells. SMGE organoids contain KRT5 (green) and ACTA2 (magenta) positive cells. Scale bars: 50 µm. **(e)** High-magnification images of each organoid type: PCCE organoid markers include BPIFA1, aTub, and KRT14 (detected by IF analysis) and *Reg3g* (detected by ISH). SSE organoid markers include LGALS7, KRT13, and SOX9 (detected by IF). SMGE organoid markers include KRT5 and ACTA2 (detected by IF). Scale bars: 25 µm. *n* = 3. SMG, submucosal gland; SPG, supraglottis; VF, vocal folds; SBG, subglottis; PCCE, pseudostratified ciliated columnar epithelium; SSE, stratified squamous epithelium; SMGE, submucosal gland epithelium; aTub, acetylated alpha-tubulin; Ap, apical side; Bs, basal side; DAPI, 4′,6-diamidino-2-phenylindole.

### Characterization of fibroblast clusters

Fibroblasts were classified into six clusters (Fig. 6a). To characterize each fibroblast cluster, Gene Ontology (GO) analysis was performed using the top 100 DEGs from Fibroblasts 1–5. In all fibroblast clusters, the top three GO terms were GO:0030198 (extracellular matrix organization), GO:0045229 (external encapsulating structure organization), and GO:0043062 (extracellular structure organization); therefore, the results for the fourth to eighth GO terms are presented in Table 1. Fibroblasts 1 and 4 included GO:0051216 (cartilage development), and Fibroblast 4 additionally contained GO:0002062 (chondrocyte differentiation).

**Fig. 6.**
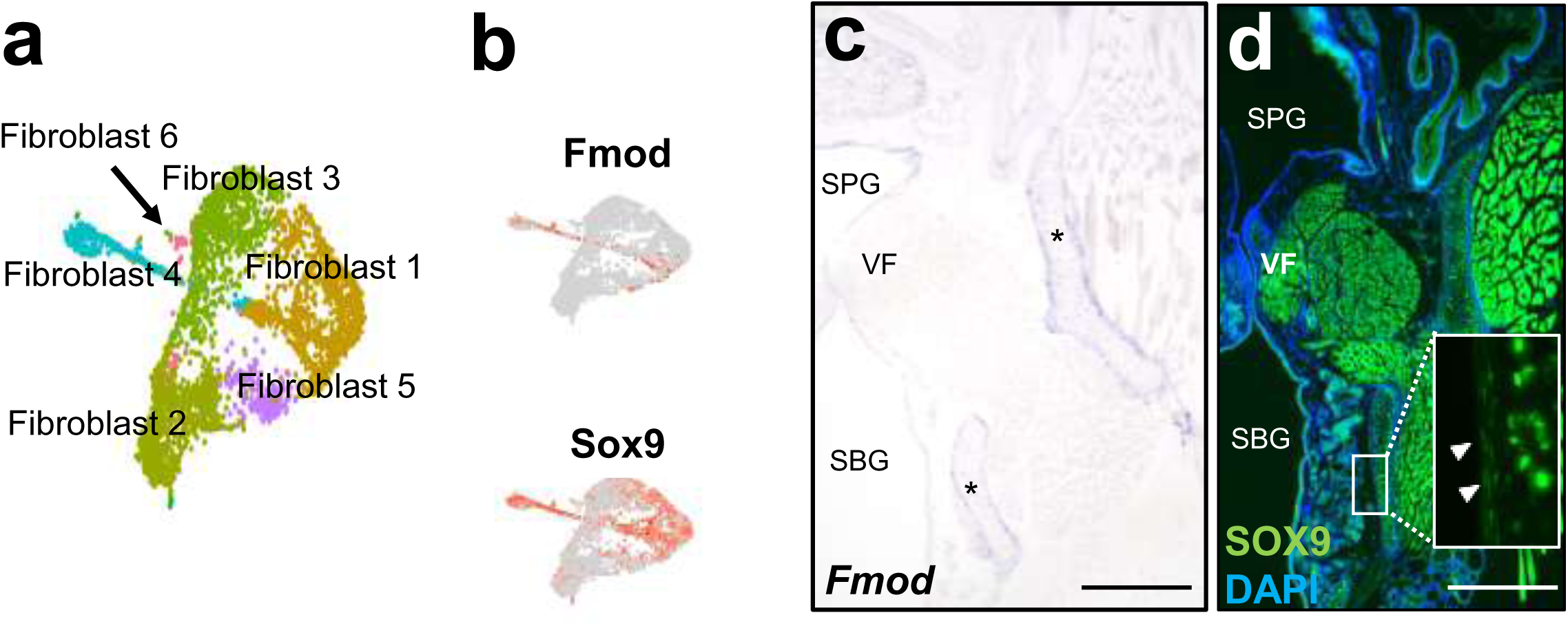
Characterization of cell types in fibroblast clusters. **(a)** Identification of six fibroblast clusters. **(b)** Feature plots of Fmod and SOX9 expression. **(c)** Tissue distribution of Fmod-positive cells. **(d)** Tissue distribution of SOX9-positive cells in mesenchymal tissue in coronal sections of the mouse larynx. White arrowheads indicate SOX9-positive perichondrial cells. Scale bars: 500 µm. *n* = 3.

Therefore, the distribution of DEGs of these clusters was examined in the feature plot. Fmod-positive cells are distributed in Fibroblast 4 and Sox9-positive cells are distributed in Fibroblast 1 and 4 (Fig. 6b). ISH (Fig. 6c) and IF (Fig. 6d) confirmed the expression of Fmod and SOX9 in perichondrial cells, suggesting that Fibroblast 4 represents perichondrial cells.

### Identification of *Lgr5*-positive fibroblast cells in lamina propria of the VFs

Spatial transcriptome analysis using PIC was conducted on axial VF sections (Fig. 7a), focusing on the anterior area (Ant), posterior area (Pst), and mesenchymal tissue between Ant and Pst (Mid) of the lamina propria, which have been reported as VF stem cell niches in human (Fig. 7a). Although statistical significance was not observed in normalized count data (counts per million [CPM] mapped reads), *Lgr5*, a known stem cell marker, exhibited higher expression in Mid (3.1 [3.7–0.9]) compared to Ant (0.8 [2.1–0]) and Pst (1.5 [1.9–0]) (Fig. 7a graph). The distribution of *Lgr5*-positive fibroblasts was further examined using ISH with RNA Scope (Fig. 7b–j). Results revealed significantly more *Lgr5*-positive fibroblasts in Ant (young: 12.7% [20.2%–7.3%], old: 11.5% [14.7%–2.8%]) and Mid (young: 18.5% [26.1%–7.7%], old: 13.4% [22.6%–4.3%]) compared to Pst (young: 3.4% [10.4%–0%], old: 3.6% [10%–0%]) (Fig. 7b). Representative images of the counted areas were shown in Fig. 7c–j. These findings suggest that *Lgr5*-positive fibroblasts are distributed throughout the lamina propria of the VFs.

**Fig. 7.**
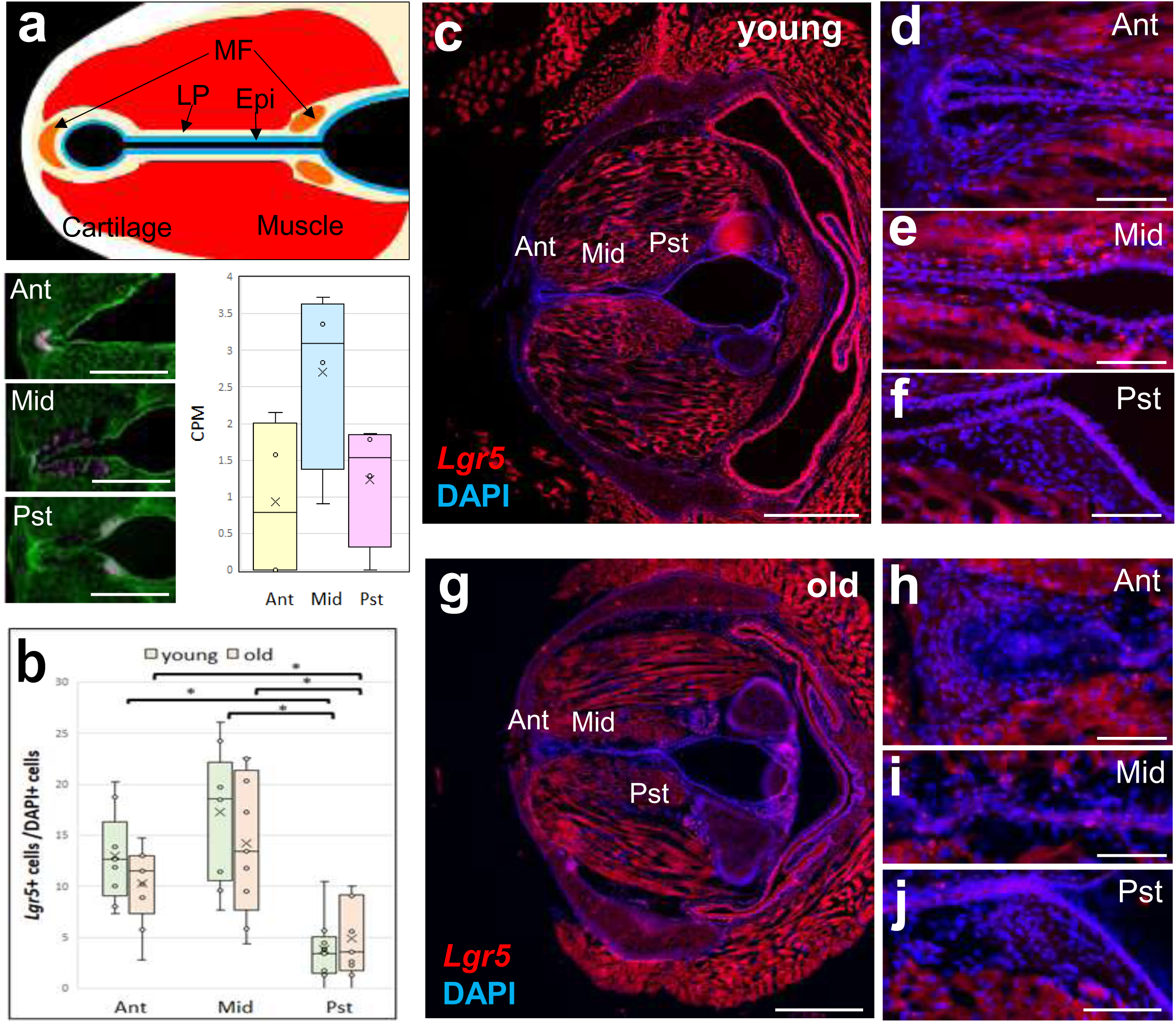
Identification of *Lgr5*-positive fibroblast cells in the mesenchymal tissue in VFs. Diagram of axial section of the larynx, showing MF (area corresponding to the human MF, orange), LP (light yellow), Epi (blue), Muscle (red), and Cartilage (white) (**a** top). Axial section images of VFs labeled with nuclei (green) and UV-irradiated areas (magenta) are shown (**a** bottom left). Scale bar: 500 µm. *n* = 4. Graph for comparison of CPM of each area **(a** bottom right). **(b-j)** Tissue distribution of *Lgr5*-positive fibroblasts in the VF lamina propria, analyzed by RNAscope. (**b)** Comparative analysis of the ratio of *Lgr5*-positive fibroblasts to DAPI-positive cells in each VF region of young and old adult mice. Asterisks indicate significant differences (*p* < 0.05, Mann–Whitney U test for young *vs.* old: Wilcoxon signed-rank sum test for comparisons among three areas). (c, g) Low-magnification images of axial sections of the VFs in young (**c**) and old (**g**) adult mice. Scale bars: 500 µm. (d–f, h–j) High-magnification images of the anterior (**d**, **h**), middle (**e**, **i**), and posterior (**f, j**) areas of the lamina propria in young and old adult mice. Scale bars: 75 µm. *n* = 6. MF, macula flava; LP, lamina propria; Epi, Epithelium; CPM, counts per million mapped reads; Ant, anterior area; Mid, middle area; Pst, posterior area.

## Discussion

In this study, we performed scRNA-seq and spatial transcriptomic analysis, gaining new insights into the cellular composition and stem cells of the larynx.

As a result of scRNA-seq, 14 epithelial cell clusters were identified (Fig. 1c–e). Consistent with a previous report, we confirmed that the PCCE clusters in the larynx closely resemble the respiratory epithelium of the trachea and other airways (23). Next, we compared the UMAP clustering data with ISH and IF results and annotated several clusters of secretory and basal cells. Analysis of secretory cell marker localization revealed that the proportion of BPIFA1-positive cells differed between the SPG and SBG. BPIFA1 is a secretory protein involved in the innate immune response to airway diseases (24). Since another BPI family member, BPIFB1, exhibits distinct yet partially overlapping localization with BPIFA1 in the nasal and oral cavities of mice (25), it is possible that other BPI family members are expressed in laryngeal tissues instead of BPIFA1. Examination of basal cell marker localization revealed that KRT14-positive basal cells were more abundant in the luminal surface epithelia of the SPG than in the SBG. In a study on mouse tracheal PCCE, KRT5-positive and KRT14-positive basal cells were identified as tissue stem cells through lineage tracing (26, 27). Under normal conditions, KRT14-positive basal cells account for approximately 20% of total basal cells in the mouse trachea, but this proportion increases following tracheal injury (28). Given that the SPG is more susceptible to damage than the SBG, it is likely that KRT14-positive cells are more prevalent in the SPG. These differences in gene expression may be attributed to the microenvironment in which each PCCE region exists or to their developmental origins. In future studies, by investigating the tissue localization of multiple DEGs in each secretory and basal cell cluster, it will be necessary to confirm the localization of the cells in each cluster, identify specific markers for secretory and basal cells in the SPG and SBG, and elucidate the functional differences between cell populations in each cluster.

Next, we examined the cell types in the SSE. To obtain further information, we conducted a subcluster analysis of *Krt13*-positive cells. This analysis identified four SSE clusters and three clusters expressing PCCE markers. Since SSE-like regions positive for KRT13 have been reported in the trachea (29), similar regions likely exist within the laryngeal PCCE, with the Krt13-positive PCCE-like cluster corresponding to the laryngeal squamociliary junction (LSCJ) (7), the transitional region between the SSE and PCCE regions. The SSEs of the VFs were found to consist of four distinct cell types with different gene expression profiles: outer layer (cluster 5), intermediate (cluster 2), cycling basal (cluster 6), and non-cycling basal cells (cluster 1). This finding aligns with array tomography observations along the dorsoventral axis, which indicated that SSEs compose one to at least three cell layers (Supplementary Fig. 1 and movie). In future studies, it will be necessary to confirm the distribution of each cluster within the laryngeal epithelium by identifying cluster-specific markers and analyzing their 3-dimentional localization using serial sections or light sheet microscopy.

In the basal layer of the medial VF region, TRP63, a well-known epithelial stem cell marker, is expressed (8, 30). In the tracheal epithelium, basal cells include several subpopulations with distinct gene expression patterns (31). To identify more specific markers for laryngeal SSE stem cells, we examined a feature plot of genes potentially involved in stemness (Supplementary Fig. 2). We found *Sox9*-positive cells in the left side of the cycling basal cluster and part of the SSBC cluster with these *Sox9*-positive cells included within *Trp63*-positive cells. SOX9 is a known stem cell marker in the SMG of the trachea (32) and in the lung (33). In human skin epidermis, SOX9 expression is primarily observed in the basal layer, and its levels increase in skin diseases such as basal cell carcinoma and squamous cell carcinoma (34). SOX9 has been reported to promote keratinization and inhibit apoptosis in keratinocytes (35). However, SOX9 expression and function in VF SSEs have not been previously reported. Consistent with the feature plot, IF analysis using anti-SOX9 confirmed SOX9-positive cells in the Trp63-positive basal cell layer, with some of these cells also positive for EdU labeling. IF analysis of laryngeal coronal sections revealed that SOX9-positive basal cells were distributed near the border between the medial and superior lateral regions of the VF (Fig. 4b). We then examined age-related differences in SOX9-positive cell numbers in this region. IF analysis of serial axial sections showed that both SOX9-positive cells and SOX9 /EdU double-positive cells were more abundant in young adult mice. SOX9 has been reported to delay aging in keratinocytes (36), suggesting that it may play a role in VF aging. Previous studies on *Sox9* in other tissues have demonstrated its function as a stem cell-related gene (37–39). SOX9 plays a critical role in regulating growth and differentiation decisions in corneal epithelia (40) and determines the outcomes of acute kidney injury, influencing whether tissue damage is temporary or progresses to fibrosis (41). Because the superior lateral to medial VF region protrudes, it is more exposed to external stimuli and foreign substances, making it more susceptible to damage. Sox9-positive cells in the laryngeal mucosa may serve a protective or regenerative function similar to SOX9-positive cells in these tissues. In future studies, it will be necessary to investigate the function of SOX9-positive cells in the vocal fold epithelium by analyzing damage model using Sox9 conditional KO mice and mice expressing fluorescent protein-fused Sox9, as well as by lineage tracing analysis using Cre-loxP system.

To establish an *in vitro* model for mechanism analysis and reagent assays, we generated organoids from laryngeal epithelia. While organoid formation from the tracheal PCCE has been widely reported (27, 42), and there are studies on SSE organoids (43) and SMG duct cell organoids (37), no studies have reported the generation of laryngeal epithelial organoids (42). Three morphological types of organoids were formed from all regions of the laryngeal mucosal epithelium in the SPG, VF, and SBG. Expression analysis confirmed that these organoids contained three epithelial cell types (PCCE, SSE, and SMGE), indicating that three types of organoids are formed under the same culture conditions. Although it was not possible to identify which marker genes/proteins were positive for each type of organoid from the IF of the sections, assuming that the Hollow type corresponds to PCCE organoid, the Solid type to SSE organoid, and the Tubular type to SMG organoid, the proportion of organoid types formed from each laryngeal epithelium region was similar to the proportions of epithelial types in the mouse larynx (Fig. 5c). The SSE organoid appeared to share properties with the SSE of VFs, as SOX9-positive cells were localized to the basal side, similar to mouse vocal folds (Fig. 5e). This suggests that these organoids could be used to study SOX9-positive basal cell functions. Although differences in BPIFA1-positive cell and KRT14-positive cell numbers were observed between SPG and SBG *in vivo*, these trends were not reflected in the organoids (Fig. 5e). This suggests that stem cells in the laryngeal PCCE of the SPG and SBG have similar organoid-forming abilities and that their gene expression may change in response to surrounding tissue signals. A study on human lung organoids co-cultured with different fibroblast types showed that gene expression varied depending on the fibroblast type (44). In future studies, after confirming the expression of additional marker genes in these organoids, it will be necessary to investigate whether the SPG–SBG gene expression differences observed *in vivo* can be reproduced in vitro by co-culturing organoids with fibroblasts from each region. Additionally, an *in vitro* lineage tracing experiment of SOX9-positive basal cells using VF organoids is needed.

GO analysis of fibroblast clusters identified cartilage-related terms in two clusters. Although cartilage was not included in scRNA-seq analysis samples, perichondrium may have been present. Fibroblast 6 expressed epithelial, secretory, and fibroblast markers (Supplementary Fig. 2), suggesting that it represents a transitional state cell cluster rather than a true fibroblast cluster. Other clusters contained large numbers of cells, making it unlikely that any were fibroblast stem cell clusters. Stem cell niches, known as macula flava, have been reported in the lamina propria of the anterior and posterior VF regions in humans (15), but their presence in mice remains unclear. We examined the expression of the stem cell markers reported in the article (15) in the feature plot, but found no distinct localization patterns (Supplementary Fig. 4). Therefore, we performed spatial transcriptomic analysis using PIC and ISH with RNAscope, revealing that *Lgr5*-expressing fibroblasts, a known stem cell marker, were scattered throughout the lamina propria. These *Lgr5*-positive fibroblasts may be similar to ABCG2-positive fibroblasts reported by Yamashita et al. (16). Previous reports have described *Lgr5*-positive mesenchymal cells in the skin (45) and lung (46, 47), where they promote epithelial progenitor differentiation via Wnt activation (46). Similarly, laryngeal *Lgr5*-positive fibroblasts may have a comparable function. *Lgr6*-positive mesenchymal cells have also been reported in mesenchymal tissues, where they promote differentiation of bronchiolar epithelial progenitors via Wnt-Fgf10 cooperation (46). In our data, *Lgr6* expresses in epithelial cell clusters extensively and in fibroblast cluster4, which is probably perichondrial cells (Supplementary Fig. 4). Only a few *Lgr6*-positive fibroblasts were detected in other fibroblast clusters, suggesting that *Lgr6*-positive cells may have a distinct role in tissue maintenance compared to *Lgr5*-positive cells. Future studies should explore the role of *Lgr5*-positive fibroblasts in injury models and co-culture experiments with laryngeal organoids treated with various factors (e.g., Wnt pathway-related factors). Additionally, we need to investigate interactions between SOX9-positive basal cells and *Lgr5*-positive fibroblasts and analyze *Lgr6* expression in the lamina propria of VFs.

However, this research includes several limitations. Insufficient annotation of epithelial clusters: Only a few known marker genes were examined. Future studies must identify DEGs with cluster-specific expression and analyze their function and localization. In addition, the larynx has a complex three-dimensional structure, but we examined only specific coronal and axial sections, except for some experiments using serial sections. Future research should use 3D imaging techniques such as tissue transparency technology to confirm the spatial distribution of each cell cluster. Lack of convergence between SOX9-positive basal cells and *Lgr5*-positive fibroblasts. The PCCE, SSE, and LSCJ exhibit distinct gene expression profiles, resulting in different distributions on UMAP. Since fibroblast clusters include cells adjacent to cartilage, muscle, SMG, and precursor cells, they may also separate into distinct clusters. Both Sox9 (37–39) and *Lgr5* (46, 48) function as stem cell markers in various organs and tissues, but their expression may be restricted to the most stem-like cells within each cluster, preventing convergence into a single cluster. Optimization of organoid generation methods: In this study, we used a single airway organoid culture condition for 14 days, generating three morphological organoid types with three types of epithelial gene expression patterns. However, only a few marker genes were examined. Further research is needed to determine whether these organoids closely resemble *in vivo* tissue and whether they are suitable for drug assays and mechanistic studies. Future studies should increase the reliability of validation using additional markers and confirm whether organoid morphology correlates with marker expression using 3D imaging. If necessary, culture conditions and durations should be optimized.

In conclusion, using scRNA-seq analysis of mouse laryngeal mucosa, we identified differences in KRT14-positive and BPIFA1-positive cell proportions in the PCCE of SPG and SBG, and discovered a SOX9-positive basal cell subpopulation in the SSE of the VF and *Lgr5*-positive fibroblasts in the lamina propria of the VF. We also successfully generated three types of organoids from mouse laryngeal mucosa. These findings contribute to our understanding of the larynx, spanning molecular biology to regenerative medicine.

## Methods

### Animals

Young adult C57BL/6 male mice (8-20 weeks old) and old adult mice (20-months old) were obtained from Japan SLC, Inc. The animal experimental protocol for this study was approved by the Animal Research Committee, Graduate School of Medicine, Kyoto University (Med Kyo 24517). All animals received humane care in compliance with the Guidelines for Animal Experiments of Kyoto University.

### Preparation of single-cell suspension for scRNA-seq

For single-cell isolation, the larynges of 8-week-old mice were dissected. The dorsal part of the larynx was cut open in the sagittal plane, and the laryngeal mucosae were collected. For each sample, the laryngeal mucosa from four male mice was pooled and incubated in 1 ml of 0.5 mM EDTA (Nacalai Tesque) for 30 min at 4°C. Then, 150 µl of 7.5% BSA (Thermo Fisher Scientific) was added, and the mucosae were collected by centrifugation at 400 x g for 5 min. After removing the supernatant, 1 ml of 0.4 mg/ml Liberase (Roche: 5401119001)/2,000 PU/ml Dispase II (Godo Shusei Co., Ltd.: 383-02281) solution was added, and the mucosae were treated at 37°C for 15 min while shaking at 300 rpm, followed by an additional 15 min incubation in a water bath at 37°C. Then, 150 µl of 7.5% BSA was added, and the epithelia were collected by centrifugation at 400 x g for 5 min. After removing the supernatant, 1 ml of 2.5% trypsin (Thermo Fisher Scientific) was added, and the sample was incubated at 37°C for 30 min. Then, 150 µl of 7.5% BSA was added, and the cells were filtered using a 40 µm tip filter. After centrifugation at 400 x g for 5 min, the supernatant was removed, and the cells were washed with 1 ml of 1% BSA/PBS. This cycle was repeated once more. After the final removal of the supernatant, cells were suspended in 50 µl of 1% BSA/PBS and used for library preparation.

### scRNA-seq and analysis

Three single-cell suspension samples were used for scRNA-seq analysis. Libraries were prepared according to manufacturer’s protocol using the Chromium Next GEM Single-Cell 3’ Reagent Kits v3.1 (10× Genomics) and the 10× Genomics Chromium Controller (10× Genomics). Sequencing was performed using DNBSEQ-G400 (MGI Tech Co., Ltd). Count matrices were generated using Cell Ranger (version 7.1.0; 10× Genomics). Count matrix was loaded into R (4.2.3)/ Seurat (version 4.3.0) and quality check and filtering were performed. After batch effect correction of data from three samples, dimensionality reduction for visualization was conducted using UMAP based on the first 18 principal components (PC 1:18) with a resolution of 0.65. Krt13-positive cell subgroups were identified through clustering analysis based on scaled gene expression values, using a threshold of log-normalized expression > 1.0. GO enrichment analyses of DEGs for each fibroblast cluster were performed using the clusterProfiler (v4.12.1) R package.

### Expression analysis

To obtain fixed samples, mice were deeply anesthetized and transcardially perfused with 4% paraformaldehyde (PFA), after which the larynges were dissected. The larynges were then immersed in 4% PFA overnight at 4°C and washed with 1xPBS. For non-fixed samples, mice were deeply anesthetized, and larynges were dissected after cervical dislocation. The larynges and organoids in Matrigel were embedded in Tissue-Tek optimal cutting temperature compound (Sakura Finetek Japan) and sectioned in the coronal or axial direction at a 10 µm thickness using a cryostat (CryoStar NX70, Thermo Fisher Scientific).

### Immunofluorescence

For staining of larynges and organoids, sections were permeabilized, blocked, and stained using primary and secondary antibodies (details summarized in Table 2). After treatment with Alexa-conjugated secondary antibodies in 1% BSA/PBS including Alexa 647-phalloidin and 4’,6-diamidino-2-phenylindole dihydrochloride (DAPI; DOJINDO), staining sections were embedded in fluoromount-G Anti-Fade (Southern Biotechnology Associates Inc.).

**Table 2.**
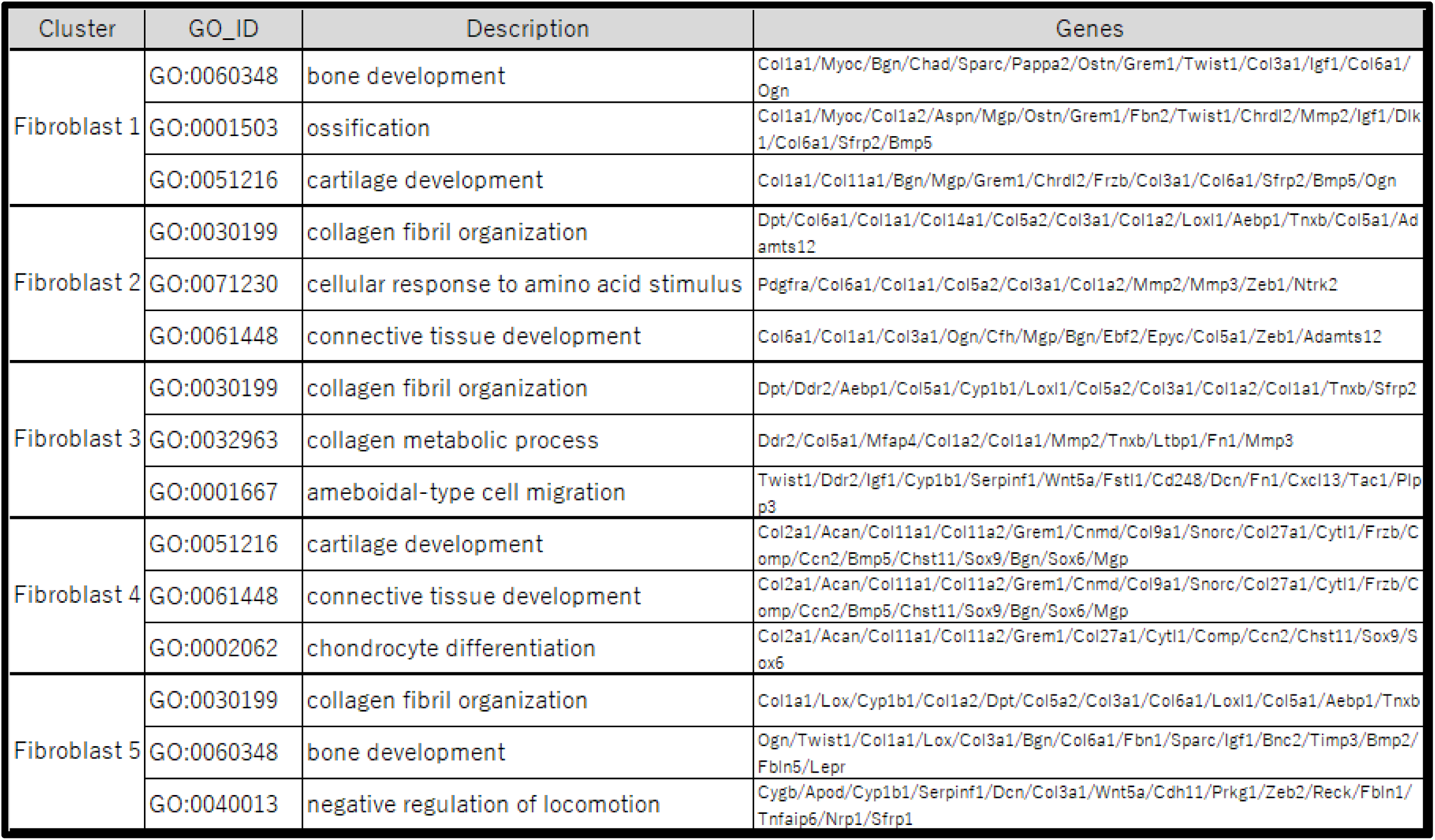
Tamura et al. GO analysis of fibroblast clusters.

### *In situ* hybridization

Fixed sections were immersed in 1× PBS for 5 min and dried at 60°C for 30 min. Slides were then fixed with 4% PFA and 0.2% glutaraldehyde in PBS at room temperature for 10 min. Sections were washed with PBST and bleached with 6% hydrogen peroxide in PBST at room temperature for 10 min, followed by another PBST wash. They were then treated with 20 µg/µl proteinase K (Roche) in PBST for 10 min, rinsed with 2 mg/ml glycine in PBST at room temperature for 10 min, and washed again with PBST. Sections were then refixed with 4% PFA and 0.2% glutaraldehyde in PBS, followed by an additional PBST wash. Next, sections were incubated in a hybridization solution at 70°C for at least 1 h, consisting of 50% formamide (Nacalai Tesque), 6 x Saline-Sodium Citrate Buffer (Nacalai Tesque; adjusted to pH 4.5 with citrate), 1% sodium dodecyl sulfate (Sigma-Aldrich), 50 µg/ml yeast RNA (Invitrogen), and 50 µg/ml heparin (Sigma-Aldrich) at 70D for at least 1 h. DIG-labeled RNA probes were hybridized with target RNA in sections in hybridization solution overnight at 70°C using a HybEZ D Oven (ACD Bio-Techne). For DIG-labeled probe preparation, DNA fragments were amplified by RT-PCR using adult mouse trachea cDNA and subcloned into a pCR-Blunt II-TOPO vector (Thermo Fisher Scientific). After digestion with appropriate restriction enzymes (New England Biolabs), DIG-labeled sense and antisense RNA probes were synthesized using a DIG RNA Labeling Kit (Roche). The RNA fragments used as probes were *Muc5b* (NM_028801, nucleotides 11054–11507), *Reg3g* (NM_011260.2, nucleotides 33–557), *Gabrp* (NM_146017.3, nucleotides 199–1657), and *Fmod* (NM_021355.4, nucleotides 1374–2211). Sections were washed with TBST and incubated in 10% sheep serum in TBST at room temperature for 1 h. They were then incubated with a 1:4000 dilution of Anti-Digoxigenin-AP, Fab fragments (Roche) at 4°C overnight. For signal detection, sections were incubated in a staining solution containing 0.25 µg/ml nitro-blue tetrazolium chloride (NBT; FUJIFILM Wako) and 0.13 µg/ml 5-bromo-4-chloro-3-indolyl phosphate (BCIP; Nacalai Tesque) in NTMT buffer, which included 100 mM NaCl (Nacalai Tesque), 0.1 M Tris-HCl (Nacalai Tesque), 10 mM MgCl_2_ (Nacalai Tesque), 0.1% Tween-20, and 0.48 g/L tetramisole hydrochloride (Sigma-Aldrich). The sections were incubated until the desired color intensity developed. IF of aTub was performed at this step. After labeling, specimens were embedded in fluoromount-G Anti-Fade.

*In situ* hybridization using the RNAscope system was performed with the RNAscope™ Multiplex Fluorescent Reagents Kit v2 with TSA Vivid Dyes (ACD Bio-Techne) according to the manufacturer’s quick guide. Briefly, sections on adhesive glass slides were immersed in 1× PBS for 5 min and dried at 60°C for 30 min. Sections were then post-fixed with 4% PFA/PBS for 1 h and dehydrated using 50% ethanol for 5 min, 70% ethanol for 5 min, and 100% ethanol for 5 min (twice). Next, sections were immersed in hydrogen peroxide for 10 min and boiled in 1× Target Retrieval Buffer for 5 min. After dehydration in 100% ethanol for 3 min, sections were incubated with Protease D at 40°C for 30 min. Sections were then hybridized with the RNAscope Target Probe (Mm-Lgr5-C3, 312171-C3; ACD Bio-Techne) at 40°C for 2 h. Post-hybridization sections were sequentially immersed in AMP1, AMP2, and AMP3 at 40°C for 15 min each. The sections were then treated with HRP-C3, TSA/Opal, and HRP blocker. Finally, sections were stained with DAPI for 30 sec and embedded in ProLong Gold Antifade Mountant (Thermo Fisher Scientific).

Specimens were observed, and images were obtained using an all-in-one fluorescent microscope (BZ-X810; KEYENCE) and BZ-X analyzer (KEYENCE).

### EdU incorporation

Cell proliferation in the larynges was assessed by detecting 5-ethynyl-2′-deoxyuridine (EdU; Thermo Fisher Scientific) incorporation into proliferating cells in tissue sections. EdU (50 mg/kg) was administered intraperitoneally to the mice, and larynges were collected 8 h post-injection, fixed, sectioned, and processed for IF. After IF, EdU was detected using the Click-iT Plus EdU Cell Proliferation Kit for Imaging Alexa 555 Dye (Thermo Fisher Scientific), according to the manufacturer’s protocol.

### Generation of laryngeal organoids

Single-cell suspensions were prepared following the same procedure as for single-cell analysis. The cell concentration was adjusted to 15,000 cells/100 µl, and the cells were mixed with 120 µl of growth factor-reduced Matrigel. The mixture was placed onto a Matrigel-coated 12-well culture insert in a 12-well culture plate, and the medium was added to the outside of the culture inserts in the 12-well culture plate. The medium for organoid formation was prepared based on a previous report (49). MTEC basic medium (Advanced DMEM-Ham’s F-12 [Thermo Fisher Scientific: 12634-010], 15 mM HEPES, 3.6 mM sodium bicarbonate, 4 mM GlultaMAX Supplement [Thermo Fisher Scientific], and 100 µg/ml Normocin [InvivoGene]); MTEC/Plus medium, which consisted of MTEC basic medium supplemented with 10 µg/ml insulin [Sigma-Aldrich], 5 µg/ml transferrin [Sigma-Aldrich], 0.1 µg/ml cholera toxin [FUJIFILM Wako], 25 ng/ml epidermal growth factor [R&D Systems, Minneapolis, MN, USA], 30 µg/ml bovine pituitary extract [Thermo Fisher Scientific], 5% FBS [Gibco: A3161001], and 0.01 µM retinoic acid [Sigma-Aldrich]; and MTEC/SF medium, which was MTEC basic medium supplemented with 5 µg/ml insulin, 5 µg/ml transferrin, 25 ng/ml cholera toxin, 5 ng/ml epidermal growth factor, 30 µg/ml bovine pituitary extract, 1 mg/ml BSA, and 0.01 nM retinoic acid. These were used as the basal medium, growth medium, and differentiation medium, respectively. Cells were cultured in MTEC/Plus medium for 7 days, after which the medium was switched to MTEC/SF medium and cultured for an additional 7 days. After 14 days, organoids embedded in Matrigel were fixed with 4% PFA for 16 h and washed with PBS for IF.

### PIC

Non-fixed axial sections of young adult laryngeal tissue were used. The PIC technique was performed as previously described (50). Briefly, tissue sections were soaked in 4% paraformaldehyde (PFA) for ten minutes, followed by permeabilization with HCl. Reverse transcription was then conducted by applying ultraviolet (UV)-responsive 6-nitropiperonyloxymethyl-caged reverse transcription primers containing a T7 promoter, unique molecular identifiers (UMIs), multiple barcodes, and a polyT sequence onto the sections. To cleave the 6-nitropiperonyloxymethyl moieties from the reverse transcription primers, the regions of interest were irradiated with UV light for three minutes using a Digital Micromirror Device (Polygon 1000-G; Mightex Systems). The total tissue lysate was collected and purified using 20 mg/mL proteinase K. Second-strand DNA synthesis was performed using the nick translation method. In *vitro* transcription reactions were conducted by transcribing synthesized cDNAs into RNAs using a T7 transcription kit. The amplified RNAs were subsequently reverse transcribed, followed by paired-end sequencing on an Illumina platform. Sequences were demultiplexed using UMI-tools based on sample barcodes and mapped to the reference genome using HISAT2. UMI-tools, HISAT2 and featureCounts were used to generate UMI count data assigned to genes. DEGs were identified using DESeq2 with a false discovery rate threshold of 0.1. In addition, DESeq2 was utilized to transform the count data into regularized log values, which were subsequently used for principal component analysis using the prcomp function in *R*.

### Electron microscopy

Mouse larynges were fixed by immersion in a fixative solution containing 0.1 M phosphate buffer with 4% PFA and 2% glutaraldehyde overnight at 4°C. The tissues were then incubated in 1% osmium tetroxide (Nacalai Tesque) for 2 h and dehydrated using ascending concentrations of ethanol. For surface structure observation, dehydrated tissues were dried by a freeze-drying method and coated with platinum-palladium. The specimens were then observed using a scanning electron microscope (SEM, JSM-7900F; JEOL). For array tomography, dehydrated tissues were embedded in epoxy resin (Luveak-812, Nacalai Tesque). Serial ultrathin sections of 400 nm thickness were collected on cleaned silicon wafer strips (approximately 5–6 mm × 10–15 mm). These sections were stained with uranyl acetate and lead citrate, and taken backscattered electron images by a SEM (JSM-7900F; JEOL) supported by Array Tomography Supporter software (version 1.6.1.0, System In Frontier Inc., Japan), which enables automated imaging (51). For 3D reconstruction, the images were stacked in order by Stacker NEO software (System In Frontier Inc.), and the resultant image stacks were processed using Dragonfly Pro software (Comet Technologies Canada Inc.).

### Statistical analysis

Data analyses were conducted using EZR (Version 1.55, Saitama Medical Center, Jichi Medical University) (52). Data are expressed as median (range, maximum–minimum value). The Mann–Whitney U test was used for PIC, RNAscope, and SOX9-positive cell analyses, while the Wilcoxon signed-rank sum test was applied to RNAscope and organoid analyses. A *p*-value <0.05 was considered statistically significant in all analyses.

### Data Availability

The datasets generated and/or analyzed during this study are available from the corresponding author upon reasonable request, and the scRNA-seq data generated in this study have been deposited in a public database.

## Acknowledgements

We thank the technical assistance in electron microscopy of Keiko Okamoto-Furuta and Haruyasu Kohda (Division of Electron Microscopic Study, Center for Anatomical Studies, Graduate School of Medicine, Kyoto University, Kyoto, Japan). Obtaining images by KEYENCE all-in-one BZ-X810 and DNA sequencing by Sanger method were performed at the Medical Research Support Center, Graduate School of Medicine, Kyoto University, which was supported by Platform for Drug Discovery, Informatics, and Structural Life Science from the Ministry of Education, Culture, Sports, Science and Technology, Japan. The process of scRNA-seq analysis, from library synthesis to obtaining fastq data from sequencing was outsourced to CyberomiX Ltd. This work was supported in part by the ISHIZUE 2023 of Kyoto University. The authors would like to thank Enago (www.enago.jp) for the English language review.

## Author contributions

K.T. contributed the acquisition, analysis, and interpretation of data, manuscript writing, and final manuscript approval; H.O. contributed conception and design, the acquisition, analysis, and interpretation of data, manuscript writing, and final manuscript approval; K.H. contributed the acquisition, analysis, and interpretation of data, manuscript writing, and final manuscript approval; S.F. contributed data analysis and final manuscript approval; T.K. contributed the acquisition, analysis, and interpretation of array tomography, and final manuscript approval; Z.Z. contributed the acquisition, analysis, and interpretation of PIC data, manuscript writing, and final manuscript approval; S.O. contributed design, analysis interpretation of PIC data, and final manuscript approval; Y.O. contributed design, analysis interpretation of PIC data, and final manuscript approval; K.O. contributed conception and design, the analysis and interpretation of data, manuscript writing, administrative support, and final manuscript approval.

## Competing interest

The authors declare no competing interests.

---

**Supplementary Fig. 1.**
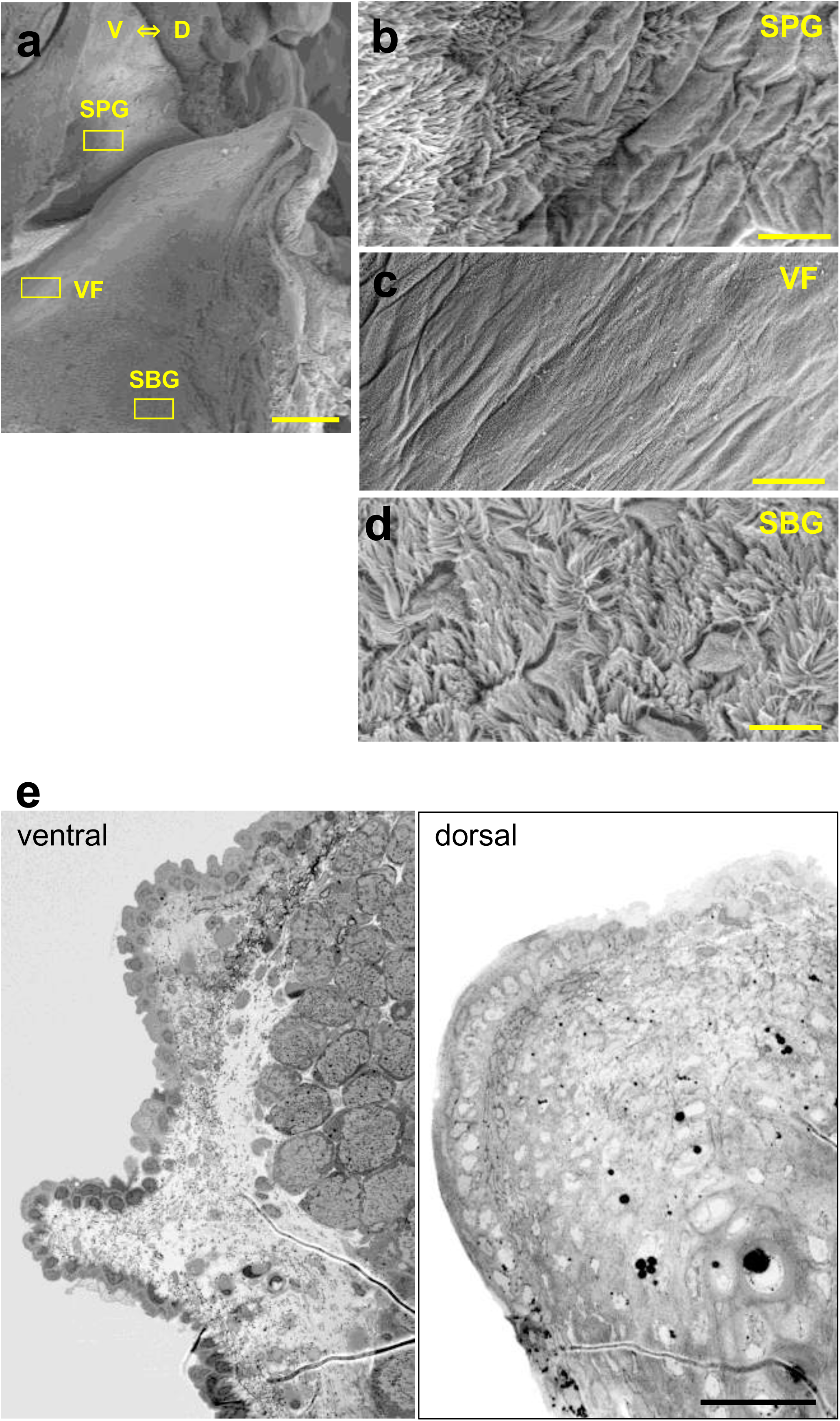
Electron microscopic analysis of the mouse larynx. Distribution of PCCE and SSE across laryngeal mucosal regions. (**a)** Sagittal section of the right half of the larynx. Scale bar: 200 µm. (b–d) High-magnification images of the laryngeal mucosal surface in the SPG (**b**), VF (**c**), and SBG (**d**). Scale bars: 10 µm. *n* = 2. (**e)** Ultrathin section images of the ventral and dorsal parts of the mouse VF. Scale bars: 40 µm. *n* = 1. SPG, supraglottis; VF, vocal fold; SBG,subglottis; SSE, stratified squamous epithelium; PCCE, pseudostratified ciliated columnar epithelium.

**Supplementary Fig. 2-1.**
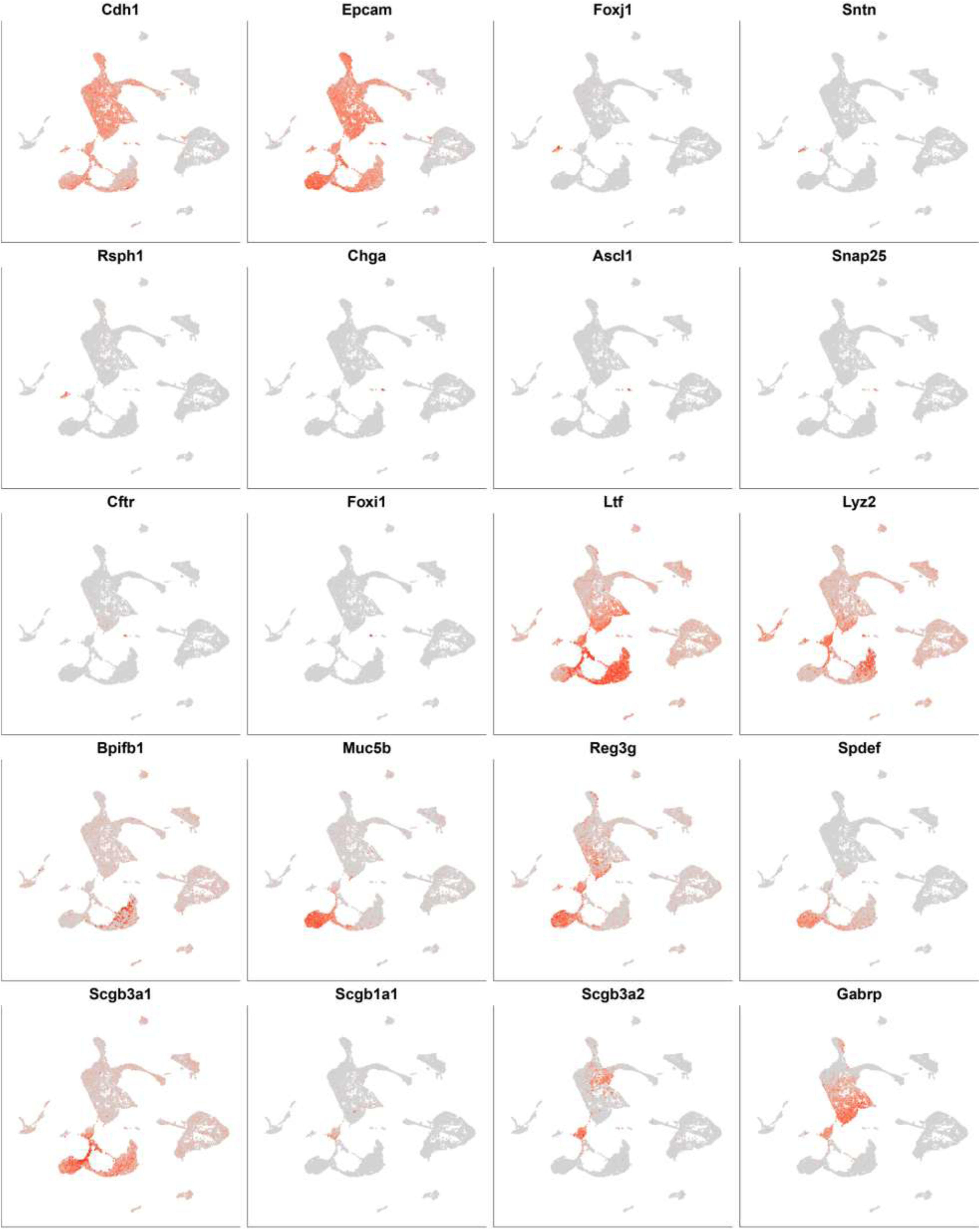
Feature plots of marker gene expression in each cluster.

**Supplementary Fig. 2-2.**
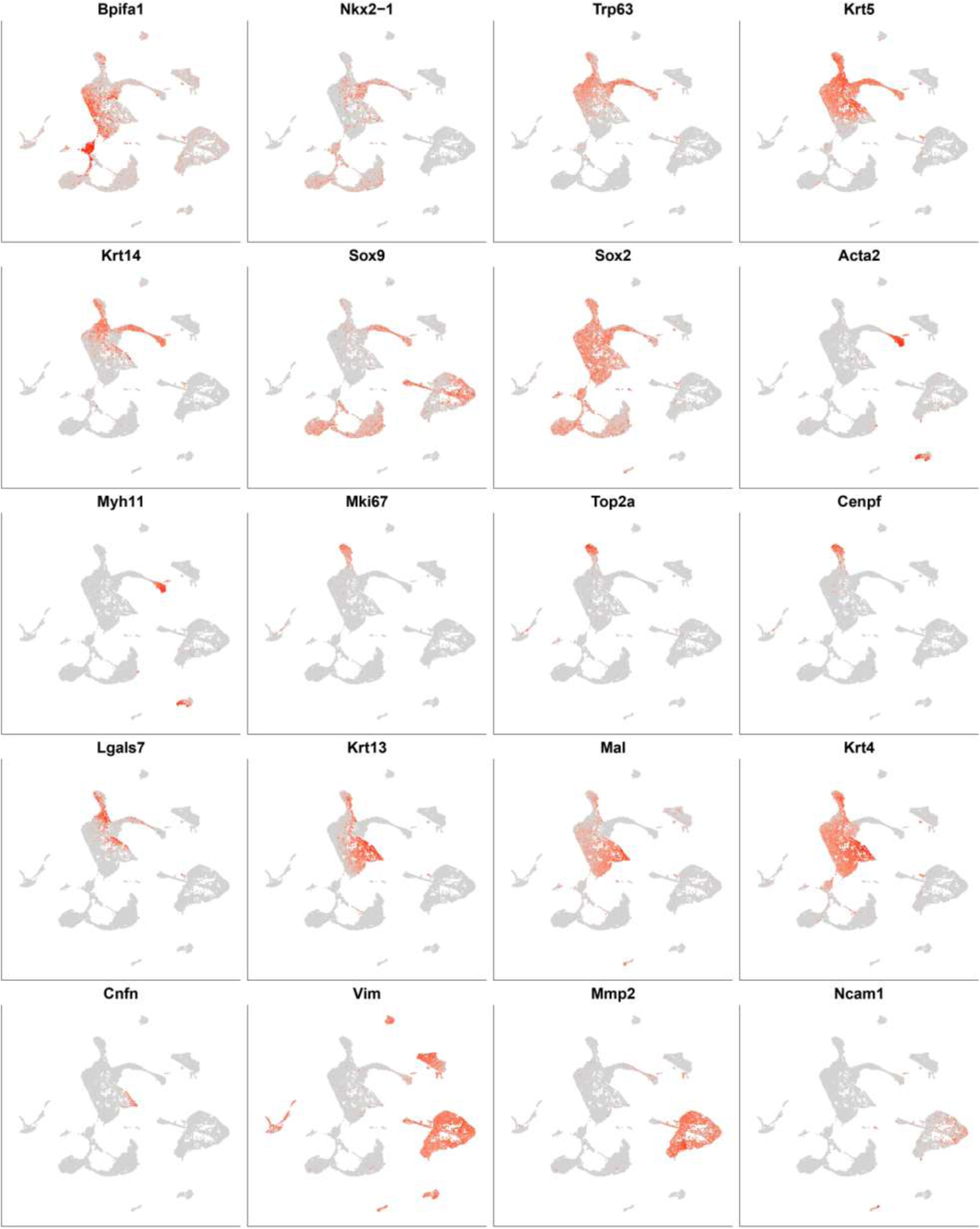
Feature plots of marker gene expression in each cluster.

**Supplementary Fig. 2-3.**
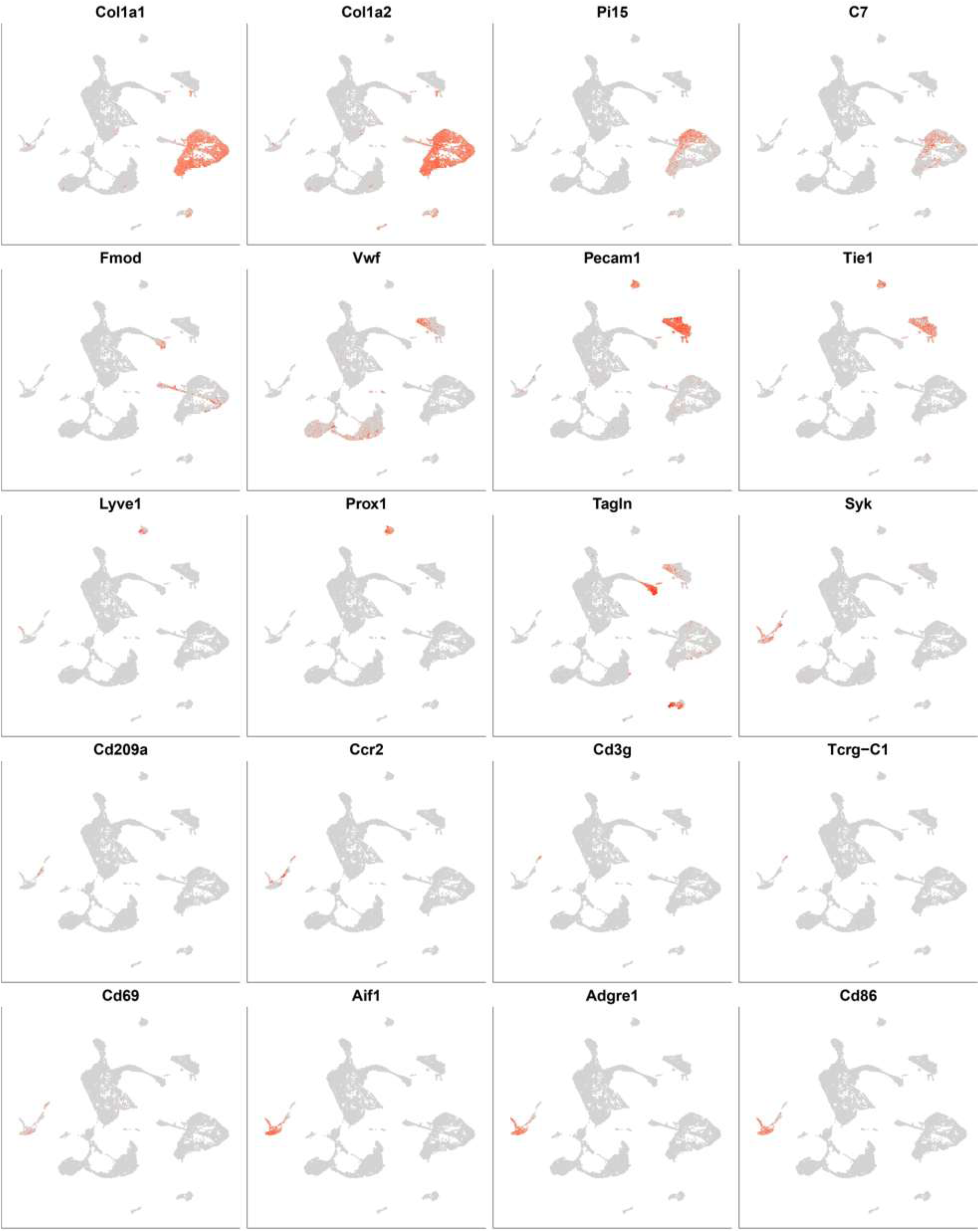
Feature plots of marker gene expression in each cluster.

**Supplementary Fig. 3.**
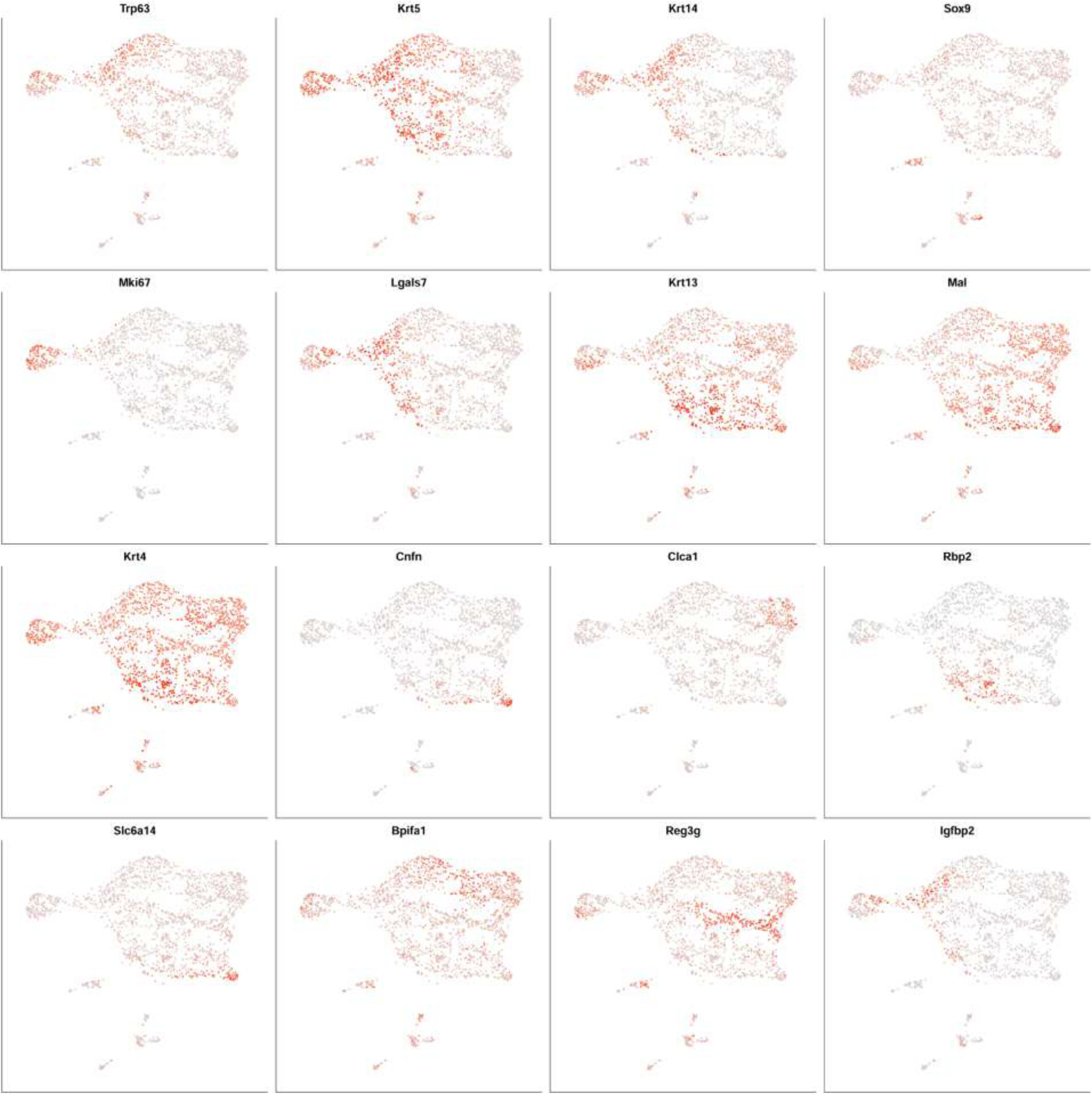
Feature plots of marker gene expression related Figure 3c.

**Supplementary Fig. 4.**
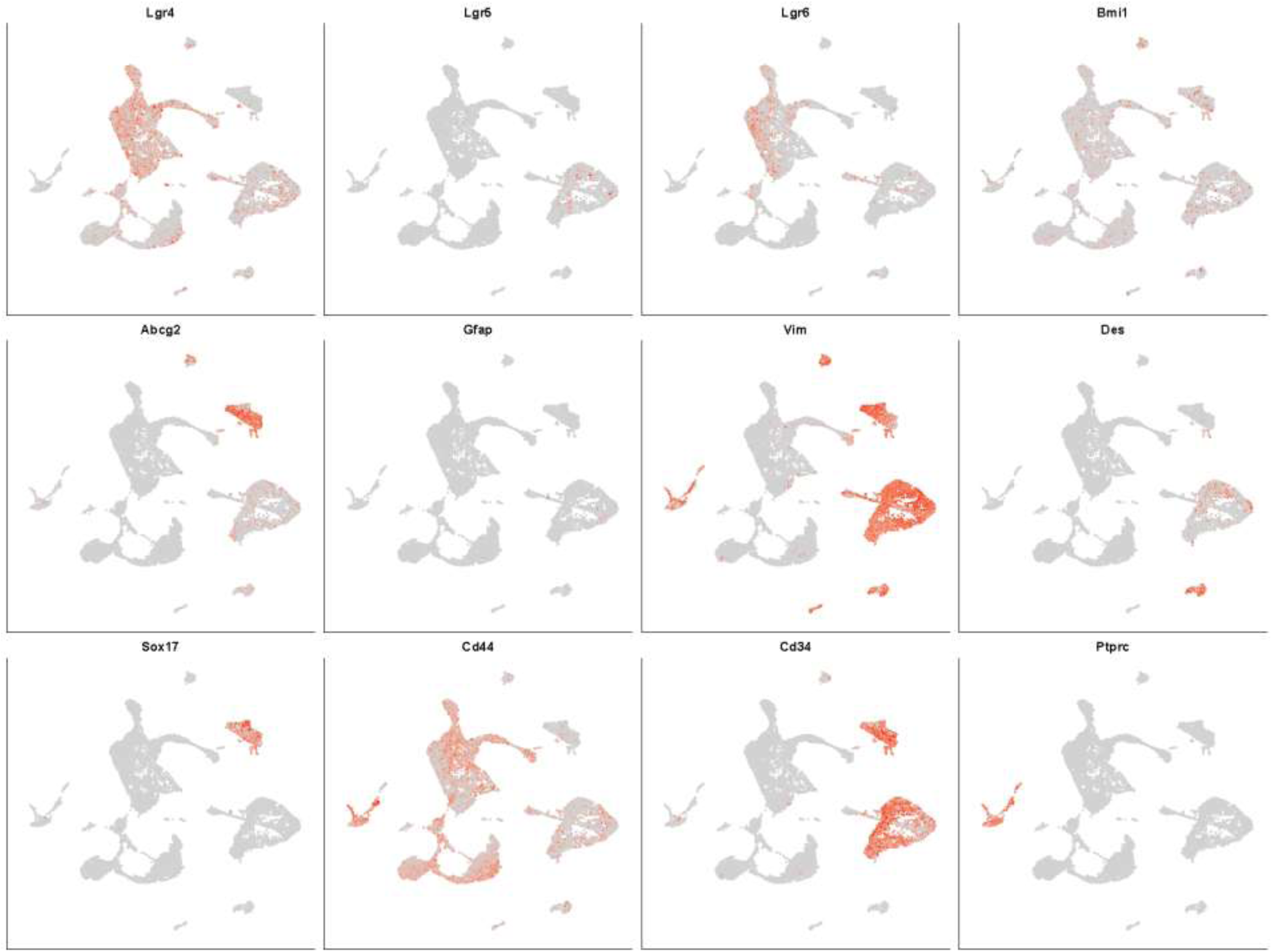
Feature plot of vocal fold stemness-related marker gene expression.

**Supplementary Fig. 5.**
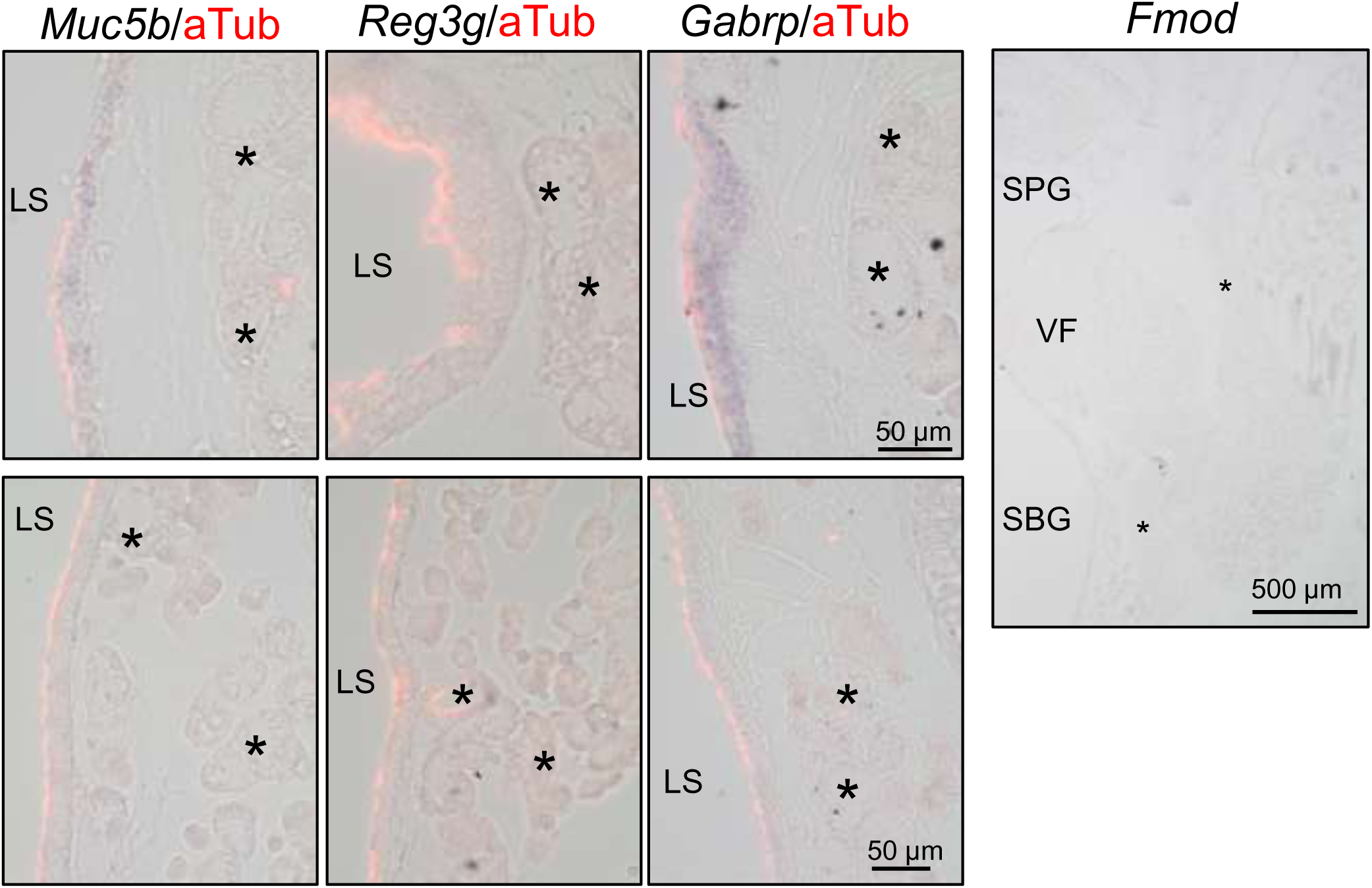
*In situ* hybridization using a sense probe as a control. *In situ* hybridization with sense probes was performed on sections adjacent to those analyzed in Figure 2c (Scale bars: 50 µm, *n* = 3) and Figure 6c (Scale bar: 500 µm, *n* = 3) as control data. Asterisks and LS indicate the SMG and luminal surface, respectively (*Muc5b*, *Reg3g,* and *Gabrp*). Labels marked by acetylated alpha-tubulin (aTub) indicate the PCCE region. Asterisks also indicate cartilage (*Fmod*). SPG, supraglottis; VF, vocal folds; SBG, subglottis; LS, luminal surface; SMG, submucosal gland; aTub, acetylated alpha-tubulin; PCCE, pseudostratified ciliated columnar epithelium

## Notes

### Competing Interest Statement

The authors have declared no competing interest.

